# A multi-agent molecular optimization framework leads to a rapid-recovery intravenous anesthetic candidate with an improved safety margin

**DOI:** 10.64898/2026.08.17.745149

**Authors:** Zhe Xue, Wei Ai, Zhuang Miao, Jinhui Sun, Zilin Wang, Wencheng Liu, Yifan Shi, Jiancheng Lv, Xianggen Liu, Bowen Ke

## Abstract

Lead optimization, the systematic refinement of therapeutic compounds through iterative structural modification, faces a dual challenge in modern drug discovery: navigating astronomically vast molecular design spaces while balancing conflicting demands on potency, pharmacokinetics, and safety. We present MASCOT (Multi-Agent SearCh for molecular OpTimization), a role-specialized multi-agent framework for molecular optimization. Integrated with a chemically constrained graph-editing search, MASCOT coordinates three specialized agents: a trade-off agent that reprioritizes competing objectives, a strategy agent that adapts how molecular edits are proposed, and a reflection agent that distills lessons from previous decisions. Computational experiments showed that MASCOT achieved the best performance over competing methods on six benchmark settings. On the SARS-CoV-2 main protease task, its mean docking-score improvement was 3.6 times that of the strongest baseline. Applied to the clinically used anesthetic remimazolam (RM), MASCOT prioritized RM-1, which showed a shorter liver microsomal half-life, higher brain exposure, and a larger therapeutic index than RM. Subsequent derivative design yielded RM-7. Extensive animal studies established RM-7 as a rapid-recovery intravenous anesthetic candidate with greater potency, faster functional recovery, a wider safety margin, and preserved flumazenil reversibility. These results demonstrate that multi-agent coordination can link adaptive molecular search to medicinal chemistry and experimental pharmacology.

**Significance Statement:** Lead optimization requires balancing potency, safety, pharmacokinetics, and ease of synthesis, because improving one property often compromises another. Managing these trade-offs while searching for better molecules is a central bottleneck in drug design. We show that a team of specialized AI agents can steer this search. Rather than generating final structures, the agents decide which goals to prioritize, how molecular edits should be proposed, and what should be learned from earlier attempts, while chemical rules ensure that every step remains valid. Applied to an approved intravenous anesthetic, this approach led to a candidate with faster recovery from anesthesia and a wider safety margin in animals. This study shows how coordinated AI agents can support practical drug discovery.

## 1. Introduction

Lead optimization is a central multi-objective challenge in drug discovery. Lead compounds must simultaneously satisfy stringent and often conflicting requirements for potency, pharmacokinetic properties, safety and synthetic accessibility (1). These coupled objectives turn molecular design into an iterative search process over a vast drug-like chemical space (2), in which improving one property can compromise others. The challenge is therefore not merely to generate candidates with favorable predicted profiles, but also to determine how search effort should be allocated among objectives as intermediate evaluations reveal new bottlenecks and trade-offs.

Computational methods have made multi-objective molecular optimization increasingly tractable (3, 4). Heuristic and evolutionary algorithms can traverse chemical space through local structural edits, but often rely on handcrafted rules, fixed proposal mechanisms or task-specific tuning (5–7). Reinforcement learning can optimize composite objectives; however, when objectives are collapsed into a fixed scalar reward, the search may favor properties that are easier to improve or exploit imperfections in the reward specification (1, 8). Deep generative models can learn molecular distributions enriched for desirable properties, but often provide limited stepwise control over chemically meaningful edits, particularly when scaffold or similarity constraints must be maintained (9–12). LLM-based agents have been applied to molecular editing, synthesis planning, tool use and workflow automation (13–17). Yet these systems have primarily emphasized candidate generation or the execution of predefined operations. What remains underdeveloped is an adaptive control architecture that can reprioritize competing objectives, revise molecular-editing strategies and learn from prior search outcomes.

The need for such control becomes acute as a multi-objective search unfolds because individual properties rarely improve in lockstep. Some may improve rapidly and then plateau, whereas others remain below target, and gains in one can erode gains in another. An effective optimizer must therefore decide when to redirect effort toward lagging objectives, when to maintain rather than further optimize objectives that have reached acceptable levels, and how to rebalance broad exploration against local refinement (8, 18, 19). We define search control as state-dependent adaptation across three dimensions: objective prioritization, edit-proposal policy and trajectory memory. Under this formulation, the controller adapts how the search is conducted, whereas molecular editing, property evaluation and constraint enforcement remain explicit and auditable computational procedures.

To instantiate this formulation, we developed MASCOT (Multi-Agent SearCh for molecular OpTimization), a role-specialized multi-agent framework for adaptive molecular search. A chemically constrained graph-editing engine performs molecular editing, property evaluation and acceptance, while three agents operate at the control layer. We evaluated MASCOT on four de novo multi-objective benchmarks and two similarity-constrained settings of a SARS-CoV-2 main protease lead-optimization benchmark. We then applied MASCOT to remimazolam (RM), a clinically used ultrashort-acting benzodiazepine anesthetic. The optimization sought to reduce dose requirements, accelerate recovery after prolonged infusion, and widen the safety margin while preserving pharmacological reversibility (20–25). With the RM core held fixed, MASCOT prioritized RM-1 for experimental follow-up. Subsequent design and experimental screening of RM-1 derivatives identified RM-7, which in animal studies showed greater potency, faster functional recovery and a wider safety margin than RM, while retaining flumazenil reversibility.

## 2. Results

### 2.1 MASCOT couples a multi-agent controller to a Monte Carlo graph-editing executor

Fig. 1 presents MASCOT as a closed-loop controller–executor architecture. At each optimization step, the executor applies local fragment addition or deletion to the current molecular graph. It removes invalid candidates, evaluates the remaining candidates with task-specific property oracles, and applies a Monte Carlo-style acceptance rule. For a molecule *x* at optimization step *t*, the composite score is written as 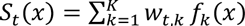, where *f* (*x*) denotes the normalized score of the objective *k*, and *w_t_*_,*k*_ is the current weight. The resulting trajectory statistics are summarized and returned to the controller.

**Fig. 1.**
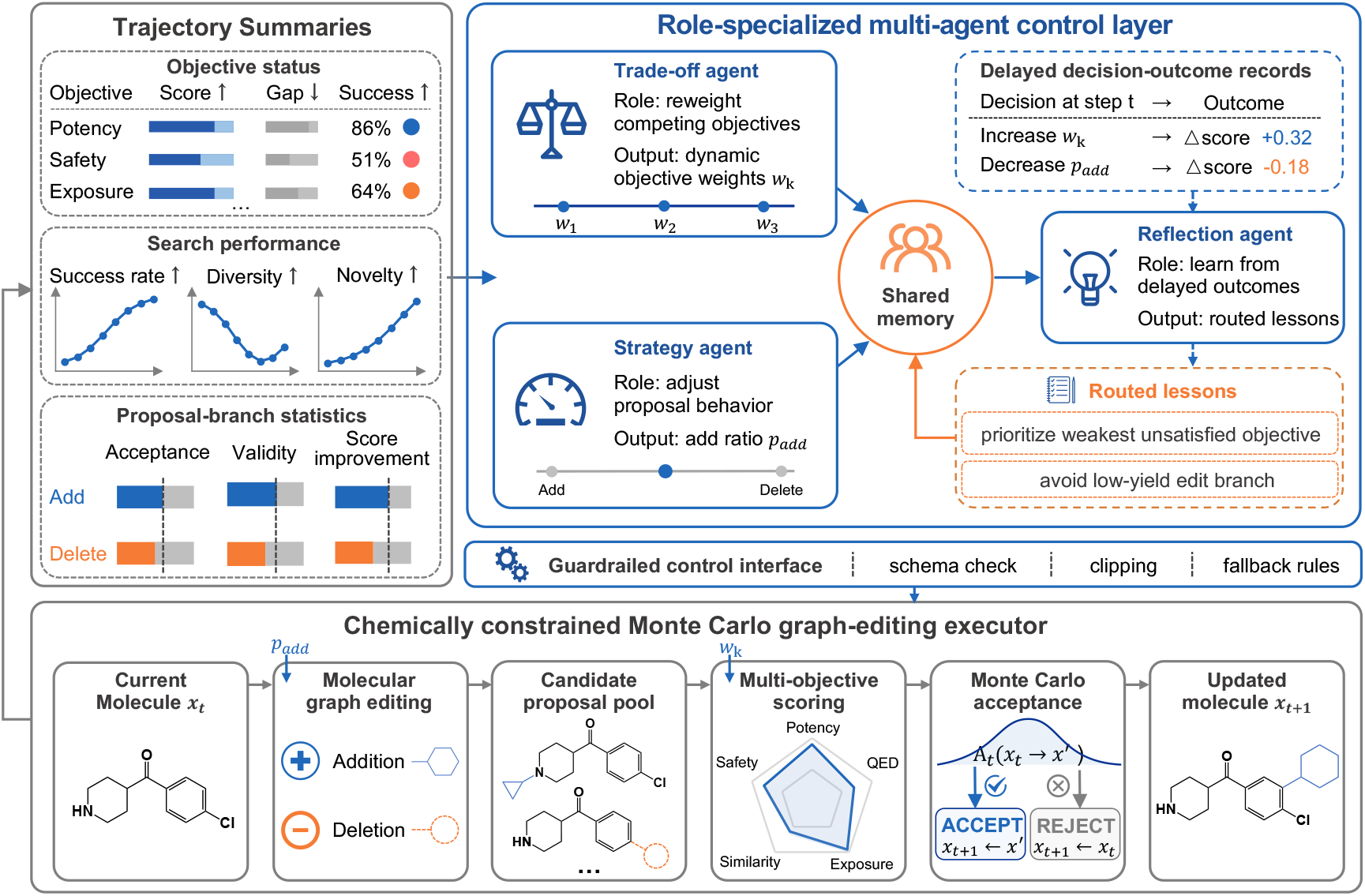
The MASCOT framework. A chemically constrained graph-editing executor proposes, scores, and accepts candidate molecules. Executor-derived trajectory summaries and delayed decision–outcome records are routed to three role-specialized agents that update objective weights, the fragment addition–deletion balance, and shared memory through a schema-validated, bounded interface. The agents control how the search proceeds but do not directly generate molecular structures.

The controller converts these observations into three bounded control outputs. The trade-off agent reallocates objective weights, the strategy agent adjusts the fragment-addition probability to regulate exploration versus refinement, and the reflection agent converts delayed decision– outcome evidence into a compact shared memory. Agent outputs pass schema validation and bound checks before being applied. Because the agents modify only search-control variables and memory, molecular editing, oracle evaluation, constraint enforcement, and acceptance decisions remain explicit and auditable computational operations.

### 2.2 Multi-agent control drives state-of-the-art multi-objective generation

We first evaluated MASCOT on de novo multi-objective molecular generation benchmarks following the MARS setting (5). The tasks require generated molecules to satisfy predicted inhibition thresholds for glycogen synthase kinase-3*β* (GSK3*β*) and/or c-Jun N-terminal kinase-3 (JNK3), together with drug-likeness (QED) (26) and synthetic accessibility (SA) (27). This benchmark was used as the main computational test because it stresses joint success under competing objectives rather than improvement of a single property. MASCOT achieved the highest product metric among the evaluated methods on the four-objective task (PM = 0.649) while retaining high joint success, novelty, and diversity (Table 1 and Table S1). Performance remained stable across five proprietary and open-weight LLM backbones, with PM values ranging from 0.620 to 0.649, indicating that the result was not dependent on a single base model (Table S2).

**Table 1.** Computational performance on the molecular generation benchmark.

| Method | GSK3β + JNK3 + QED + SA | | GSK3β + QED + SA | | JNK3 + QED + SA | | GSK3β + JNK3 | |
| --- | --- | --- | --- | --- | --- | --- | --- | --- |
|  | SR | PM | SR | PM | SR | PM | SR | PM |
| GCPN (28) | 0.000 | 0.000 | 0.000 | 0.000 | 0.000 | 0.000 | 0.035 | 0.002 |
| JT-VAE (29) | 0.054 | 0.015 | 0.096 | 0.063 | 0.218 | 0.131 | 0.033 | 0.002 |
| RationaleRL (30) | 0.750 | 0.294 | 0.891 | 0.270 | 0.787 | 0.131 | 0.842 | <u>0.686</u> |
| GA+D (31) | 0.860 | 0.310 | 0.890 | 0.610 | 0.860 | 0.430 | 0.850 | 0.360 |
| MARS (5) | 0.923 | 0.547 | **0.995** | <u>0.680</u> | 0.913 | <u>0.674</u> | **0.995** | 0.518 |
| RetMol (12) | <u>0.969</u> | <u>0.611</u> | 0.913 | 0.521 | <u>0.951</u> | 0.578 | 0.847 | 0.436 |
| MASCOT | **0.977** | **0.649** | <u>0.993</u> | **0.720** | **0.985** | **0.680** | <u>0.991</u> | **0.707** |

SR is the fraction of molecules satisfying all thresholds; Nov is the fraction absent from the training set; Div is structural diversity; PM = SR × Nov × Div. Success thresholds for each task: QED ≥ 0.6, SA ≥ 0.67, and GSK3β /JNK3 inhibition ≥ 0.5. For each method, 5,000 molecules were generated and evaluated over 10 independent runs. RetMol was rerun using its open-source implementation; other baseline results were taken from MARS. The best values are shown in bold, and the second-best values are underlined. All main-text experiments used Gemini 2.5 Pro as the agent backbone.

MASCOT also improved optimization efficiency, reaching PM = 0.5 approximately 55 steps earlier than the no-agent control and converging to a higher final value (Fig. 2A). To determine whether this advantage arose from the control layer, we compared the full framework with five reduced controllers that shared an identical graph-editing executor (Fig. 2B). Each reduced controller captured only part of the improvement (PM = 0.588–0.618), whereas the full framework achieved the best overall balance of success, novelty, and diversity. These results support complementary roles for the trade-off, strategy, and reflection agents.

**Fig. 2.**
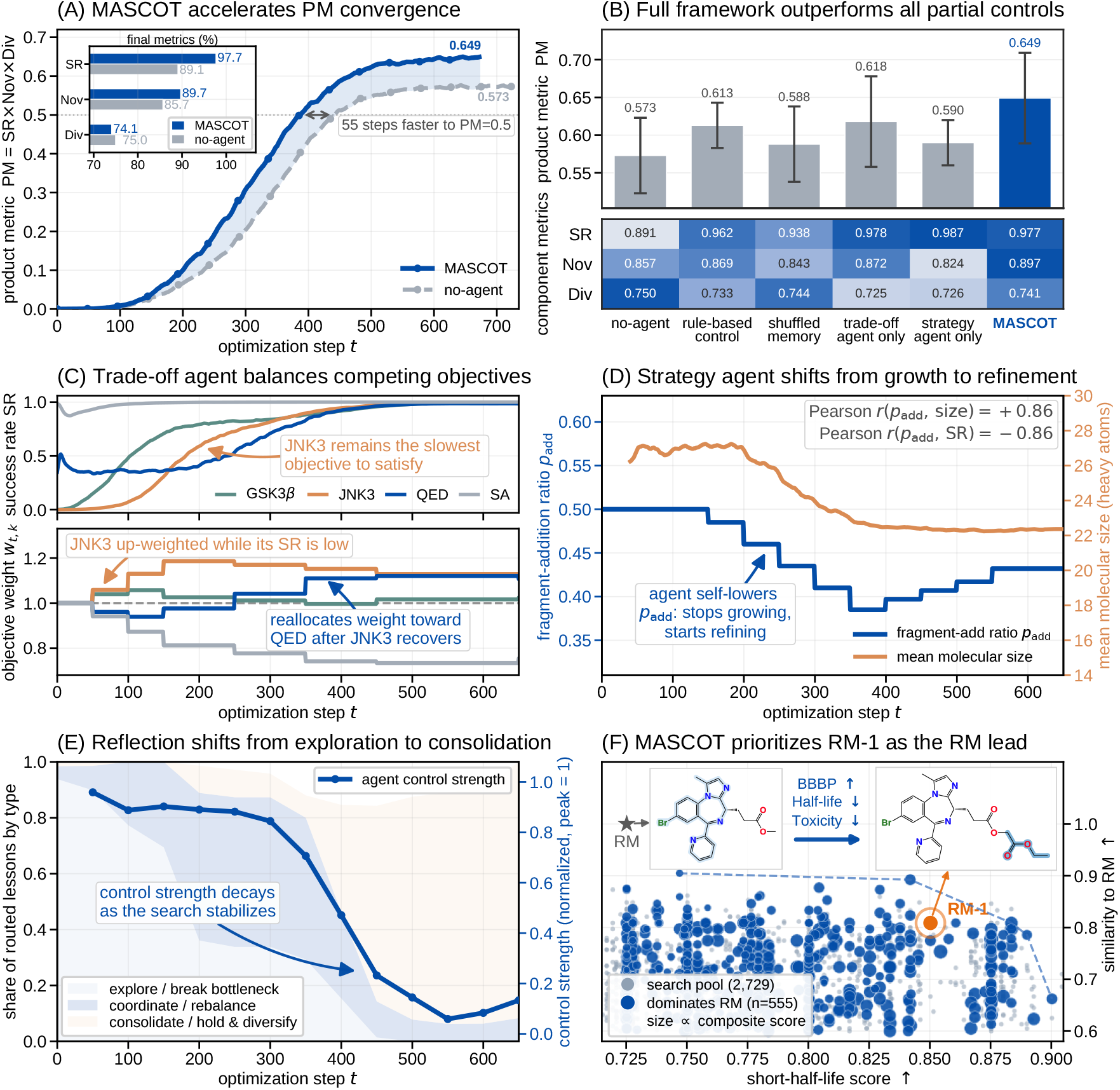
Coordinated multi-agent control drives benchmark performance (A–E) and prioritizes the remimazolam lead (F). Panels A–E use the GSK3β+JNK3+QED+SA task. (A) Product metric (PM = SR × Nov × Div) trajectories for the full framework versus a no-agent control (curves are averages over 10 runs). MASCOT reaches PM = 0.5 about 55 steps earlier and converges to a higher final value. (B) PM for the full framework and five reduced controllers (top; error bars, SD across 10 runs). The heat map shows the corresponding SR, Nov, and Div values (bottom; colors are scaled within each metric row). The rule-based variant updated variables using deterministic heuristics; the shuffled-memory variant retained the reflection-agent interface but randomly re-paired decisions with subsequent outcomes. The full framework achieved the best overall balance. (C) The trade-off agent tracks each objective’s success rate (top) and reallocates the objective weights accordingly (bottom). JNK3 remains the slowest objective to satisfy; its weight is increased while its SR is low, and weight is later reallocated toward QED after JNK3 recovers. (D) Across the trajectory, the fragment-addition ratio padd correlated positively with molecular size (Pearson r = +0.86) and negatively with SR (Pearson r = −0.86; pooled over agent decision points across 10 runs). This pattern is consistent with the agent suppressing further expansion when additional fragment growth no longer improved success. (E) Reflection agent behavior is shown on two aligned vertical axes sharing the optimization-step axis. The stacked bands show the composition of routed lesson types (left), whereas the blue curve shows overall control strength (right). Lesson routing shifts from exploration early to consolidation late, and control strength declines as the search stabilizes. (F) Remimazolam (RM) application. The strict-scaffold search comprised 100 parallel chains run for 50 steps and yielded 2,729 unique candidates. Each point represents one candidate; the 555 candidates that dominated RM on all three objectives are highlighted, and point size is proportional to the composite score. Candidates are positioned by short-half-life score (x-axis) and similarity to RM (y-axis). From this pool, MASCOT prioritized RM-1 (red), which preserved the RM scaffold while improving the predicted profile, with higher BBBP, a shorter metabolic half-life, and lower toxicity risk.

The control trajectories explain how this gain emerged. The trade-off agent redirected optimization pressure toward the lagging JNK3 objective and relaxed that pressure after its success rate recovered (Fig. 2C). Continued fragment addition became associated with larger molecules and lower success rates. The strategy agent therefore reduced the fragment-addition ratio, shifting the search from expansion to refinement (Fig. 2D). In parallel, reflection lessons shifted from early exploration and bottleneck breaking to late consolidation and diversity preservation, while intervention strength declined as the search stabilized (Fig. 2E). Together, these results indicate that MASCOT improved performance by coordinating how the search evolved.

We next tested transfer to a similarity-constrained SARS-CoV-2 main protease (Mpro, PDB ID: 7L11) lead-optimization task following the RetMol setting (12). We started from eight weak inhibitors and optimized predicted binding (32) under QED, SA, and similarity constraints. MASCOT produced the largest mean docking-score improvements among the evaluated methods (33) under both similarity thresholds (Tables S3 and S4). This indicates that the same search-control framework extends from de novo generation to constrained lead optimization.

### 2.3 Scaffold-constrained optimization of remimazolam prioritizes RM-1

We next applied MASCOT to a prospective lead-optimization problem starting from RM, a clinically used ultrashort-acting benzodiazepine anesthetic (20). Although RM has a rapid onset and short duration after bolus administration, its recovery after prolonged administration, dose requirements, and safety margin can still be improved (21, 22, 34, 35). We therefore formulated RM optimization as a scaffold-preserving multi-objective task. The RM core scaffold was held fixed, and graph edits were restricted to the side chains. The optimization objectives were a shorter predicted metabolic half-life, greater predicted blood–brain barrier penetration (BBBP), and lower predicted toxicity risk. All three oracle outputs were transformed to benefit-oriented scales, so that higher values consistently indicated more desirable predicted properties.

MASCOT identified RM-1 as the computational lead for synthesis and experimental validation (Fig. 2F). RM-1 preserved the RM scaffold while modifying the ester-containing side chain. Its predicted binding pose at the GABA_A_ receptor is shown in Fig. S1. This modification was consistent with the intended strategy of altering disposition-related properties while retaining the parent pharmacophore. Detailed search settings are provided in SI Appendix.

Experimental assays supported the computational prioritization of RM-1. In human liver microsome stability assays, RM-1 showed a shorter liver microsomal half-life than RM (6.49 min vs. 19.66 min; Table 2), indicating faster microsomal metabolism. RM-1 also showed improved anesthetic potency, with a lower median effective dose (ED_50_) than RM (23.85 vs. 38.23 mg/kg; Table 2), higher brain exposure (203.14 vs. 177.05 μM; Fig. 4E), and a larger therapeutic index (TI: 9.94 vs. 6.56; Fig. 4E).

**Table 2.**
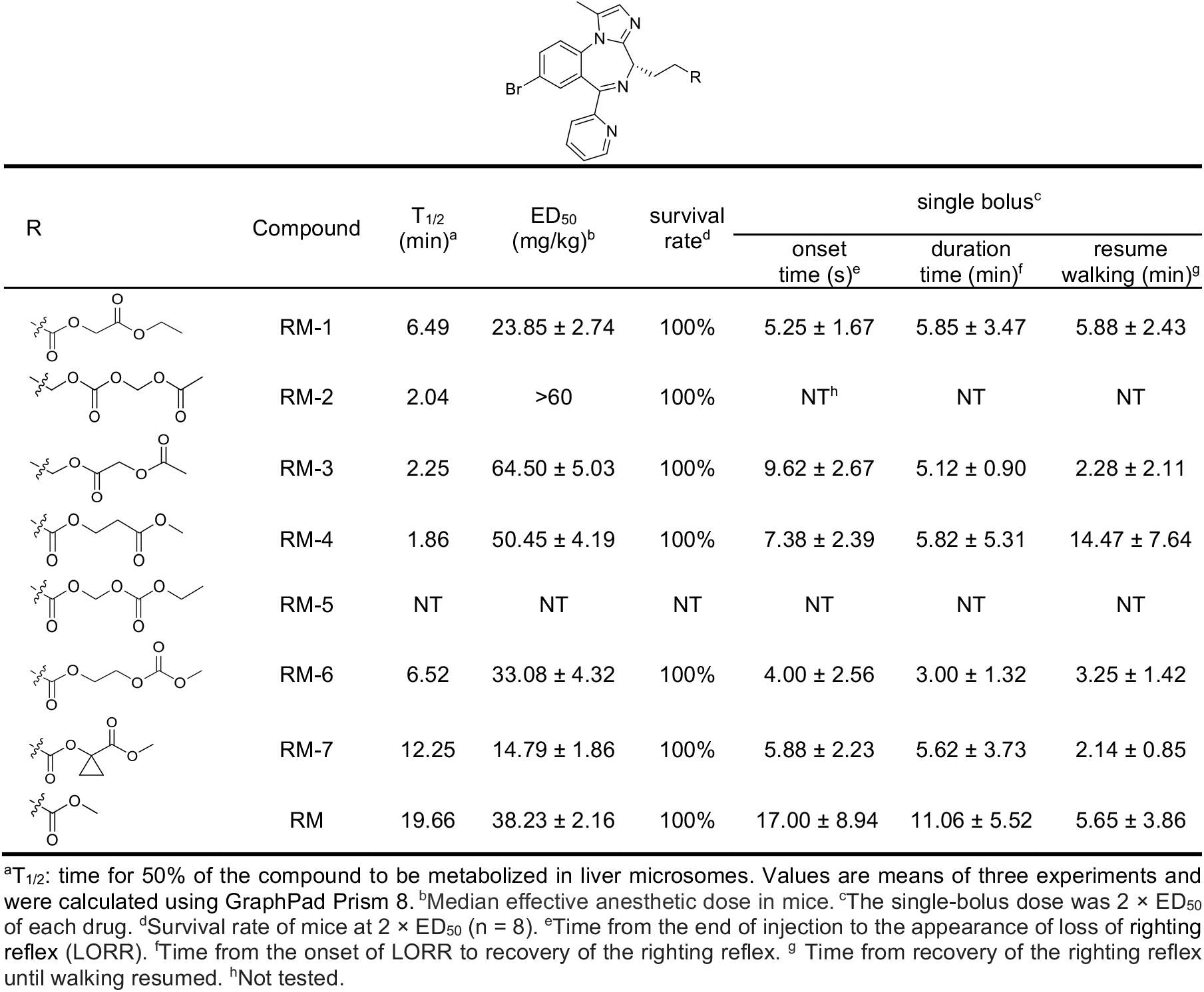
Metabolic stability and in vivo anesthetic properties of RM, RM-1, and RM-1 derivatives.

| R | Compound | T<sub>1/2</sub><br>(min)<sup>a</sup> | ED<sub>50</sub><br>(mg/kg)<sup>b</sup> | survival<br>rate<sup>d</sup> | single bolus<sup>c</sup> |  |  |
| --- | --- | --- | --- | --- | --- | --- | --- |
|  |  |  |  |  | onset<br>time (s)<sup>e</sup> | duration<br>time (min)<sup>f</sup> | resume<br>walking (min)<sup>g</sup> |
|  | RM-1 | 6.49 | 23.85 ± 2.74 | 100% | 5.25 ± 1.67 | 5.85 ± 3.47 | 5.88 ± 2.43 |
|  | RM-2 | 2.04 | >60 | 100% | NT<sup>h</sup> | NT | NT |
|  | RM-3 | 2.25 | 64.50 ± 5.03 | 100% | 9.62 ± 2.67 | 5.12 ± 0.90 | 2.28 ± 2.11 |
|  | RM-4 | 1.86 | 50.45 ± 4.19 | 100% | 7.38 ± 2.39 | 5.82 ± 5.31 | 14.47 ± 7.64 |
|  | RM-5 | NT | NT | NT | NT | NT | NT |
|  | RM-6 | 6.52 | 33.08 ± 4.32 | 100% | 4.00 ± 2.56 | 3.00 ± 1.32 | 3.25 ± 1.42 |
|  | RM-7 | 12.25 | 14.79 ± 1.86 | 100% | 5.88 ± 2.23 | 5.62 ± 3.73 | 2.14 ± 0.85 |
|  | RM | 19.66 | 38.23 ± 2.16 | 100% | 17.00 ± 8.94 | 11.06 ± 5.52 | 5.65 ± 3.86 |

<sup>a</sup>T<sub>1/2</sub>: time for 50% of the compound to be metabolized in liver microsomes. Values are means of three experiments and were calculated using GraphPad Prism 8. <sup>b</sup>Median effective anesthetic dose in mice. <sup>c</sup>The single-bolus dose was 2 × ED<sub>50</sub> of each drug. <sup>d</sup>Survival rate of mice at 2 × ED<sub>50</sub> (n = 8). <sup>e</sup>Time from the end of injection to the appearance of loss of righting reflex (LORR). <sup>f</sup>Time from the onset of LORR to recovery of the righting reflex. <sup>g</sup>Time from recovery of the righting reflex until walking resumed. <sup>h</sup>Not tested.

### 2.4 RM-7 is a rapid-recovery anesthetic candidate with a wider safety margin

Because RM-1 served only as an intermediate lead, we performed focused side-chain derivatization while retaining the RM scaffold, yielding RM-2–RM-7. Integrated screening of metabolic stability, potency, recovery, and survival narrowed the series to RM-7. RM-5 was chemically unstable (Table S5), RM-2–RM-4 were cleared too rapidly, and RM-6 had a less favorable potency–duration profile. The derivatives with the shortest half-lives were not the best anesthetics. Recovery therefore appears to depend on a balanced exposure window determined by metabolic stability, potency, and brain exposure, rather than on the shortest possible half-life.

RM-7 showed the most favorable profile among RM, RM-1, and the derivatives. Its liver microsomal half-life was intermediate (12.25 min vs. 6.49 min for RM-1 and 19.66 min for RM), avoiding the half-lives below 3 min observed for RM-2–RM-4 (Table 2). In mice, RM-7 lowered the ED_50_ to 14.79 ± 1.86 mg/kg (vs. 23.85 for RM-1 and 38.23 for RM) and, at 2 × ED_50_, shortened post-LORR walking recovery to 2.14 ± 0.85 min (vs. 5.88 and 5.65 min) while preserving rapid onset and short hypnotic duration (Table 2). In rabbits, recovery after continuous infusion (0.5–6 h) was shorter and more stable than with RM (Fig. 3B), and maintenance-dose and EEG analyses indicated comparable anesthetic depth (Fig. 3C, E, and F).

**Fig. 3.**
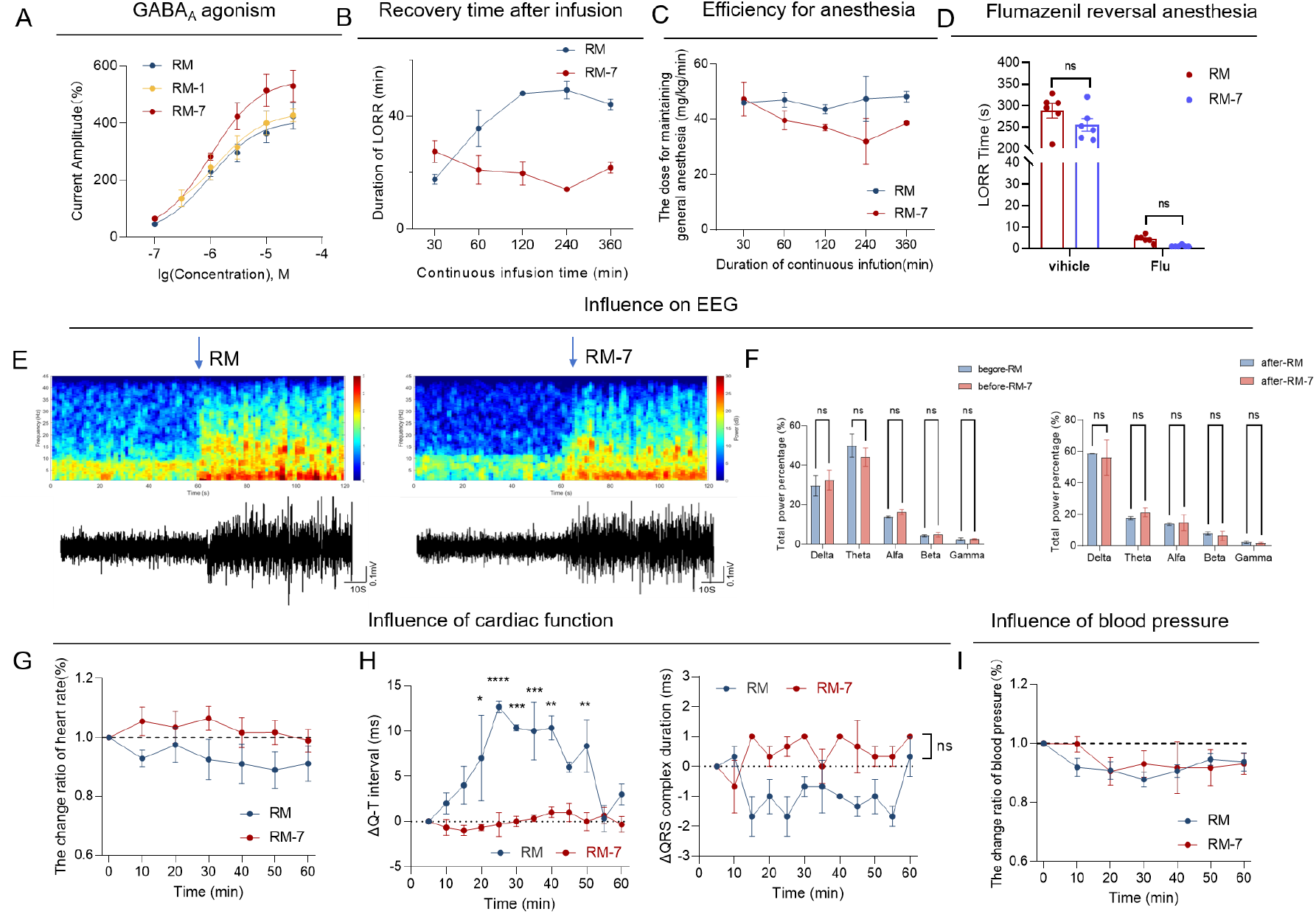
Target validation of RM-7 and physiological monitoring during continuous infusion. (A) gamma-aminobutyric acid type A (GABAA) receptor potentiation (positive allosteric modulation) in vitro (n = 3). (B) Time to recovery of the righting reflex after cessation of continuous infusion (n = 3). (C) Dose of the compounds required to maintain anesthesia during continuous infusion (n = 3). (D) Flumazenil reversal experiment (n = 3). (E) Representative EEG spectrograms and traces before and after administration of RM or RM-7 (n = 3). (F) Relative power in the delta, theta, alpha, beta, and gamma EEG bands before and after administration (n = 3). (G) Effects of the compounds on heart rate (n = 3). (H) Changes in QT interval and QRS-complex duration during continuous infusion (n = 3). (I) Relative changes in blood pressure during continuous infusion (n = 3). Data are presented as mean ± SEM. ns, P > 0.05; *P < 0.05; ***P < 0.001; ****P < 0.0001.

RM-7 retained the expected GABA_A_ receptor activity and flumazenil reversibility. In vitro, it potentiated GABA_A_ receptor currents more strongly than RM or RM-1 (EC_50_ = 1.058 μM; maximal amplitude 529.01%; Fig. 3A). In vivo, flumazenil rapidly reversed anesthesia for both RM and RM-7, reducing recovery to a few seconds with no significant between-group difference (Fig. 3D), supporting RM-7 as a more potent yet still flumazenil-reversible benzodiazepine-like anesthetic.

Safety and exposure studies further favored RM-7. During continuous infusion, RM-7 produced smaller fluctuations in heart rate and blood pressure and less QT-interval perturbation than RM (Fig. 3G–I). No major serum biochemical abnormalities, histological abnormalities in the heart, liver, or kidney, or abnormal loss of body weight were observed (Fig. 4A–C). Acute toxicity testing yielded a higher LD_50_ than RM or RM-1 (355.54 vs. 250.95 and 238.08 mg/kg) and thus a larger therapeutic index (24.04 vs. 6.56 and 9.94), alongside higher brain exposure (247.04 vs. 177.05 and 203.14 μM; Fig. 4E). Behavioral testing showed less residual impairment: open-field activity, rotarod performance, and Y-maze novel-arm exploration recovered without the delays observed after RM (Fig. 4D, F, and G). Overall, RM-7 combines stronger potency, faster functional recovery, flumazenil reversibility, improved brain exposure, fewer cardiac conduction abnormalities, and a wider safety margin.

**Fig. 4.**
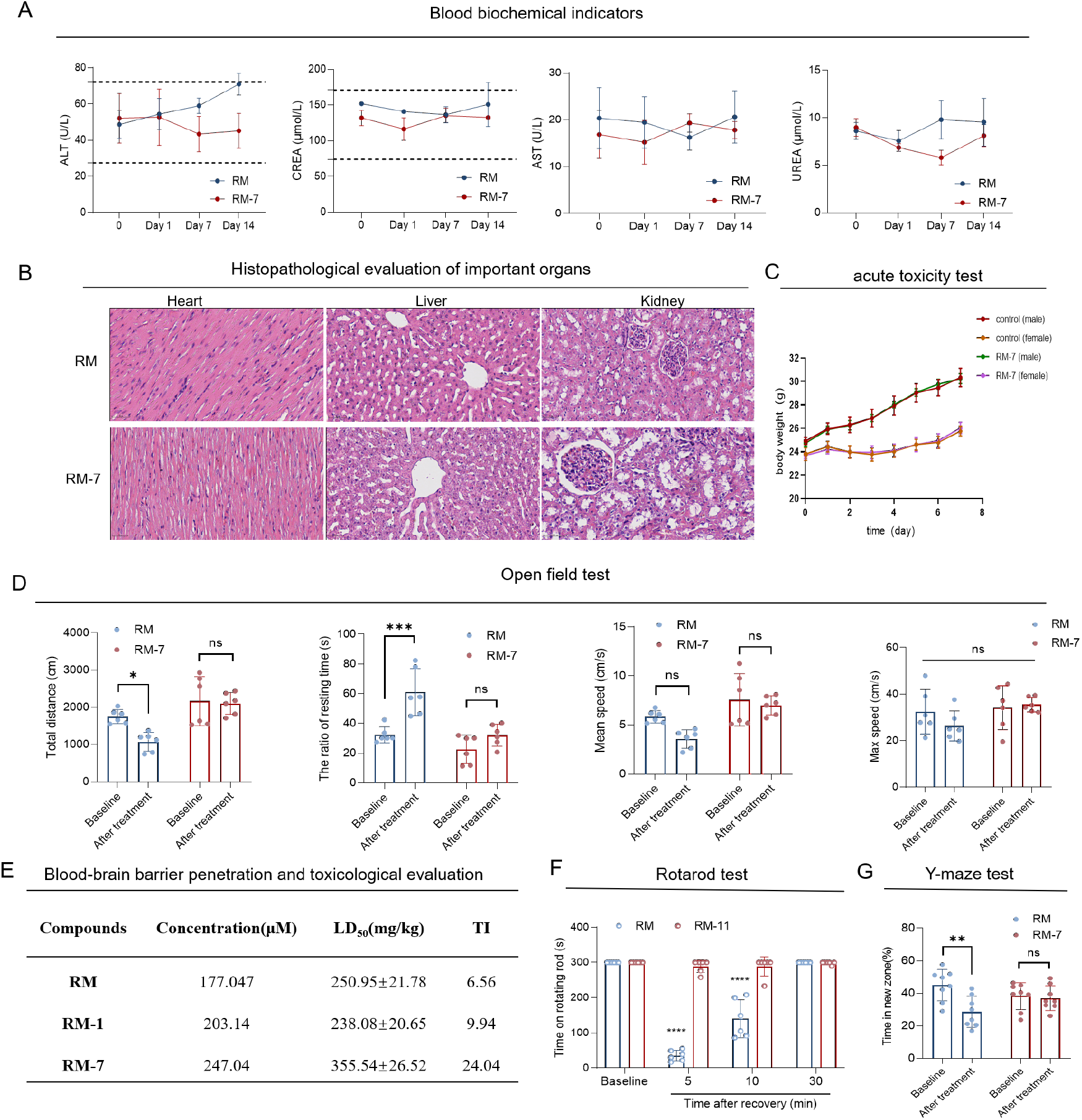
Safety assessment of RM-7 in vivo. (A) Serum alanine aminotransferase (ALT), creatinine (CREA), aspartate aminotransferase (AST), and urea levels at the indicated time points (n = 3). (B) Histopathological evaluation of major organs 7 days after continuous infusion in rabbits. Both RM and RM-7 were infused continuously for 1 h. Scale bar = 50 μm (20×) (n = 3). (C) Body-weight trajectories of male and female mice after RM-7 (5 × ED50) or control treatment (n = 8). (D) Open-field behavior before and after RM or RM-7 at 2 × ED50 (n = 8). (E) Brain exposure, LD50, and therapeutic index for RM, RM-1, and RM-7 (n = 8). (F) Rotarod performance after recovery from anesthesia in mice (n = 8). (G) Y-maze novel-arm exploration after recovery from anesthesia in mice (n = 8). Statistical analysis was performed using one-way ANOVA followed by Tukey’s post hoc tests. Data are presented as mean ± SEM. ns, P > 0.05; *P < 0.05; **P < 0.01.

## 3. Discussion

MASCOT reframes multi-objective molecular optimization as a search-control problem. Rather than generating final structures or optimizing a fixed reward, three specialized agents regulate objective pressure, proposal behavior, and trajectory memory. Molecular editing, scoring, constraint enforcement, and acceptance remain explicit computational procedures. Because all ablations used the same graph-editing executor, the superior balance of success, novelty, and diversity achieved by the full framework reflects the contribution of coordinated control. Its transfer from de novo generation to similarity-constrained SARS-CoV-2 Mpro optimization further suggests that MASCOT functions as a reusable control protocol rather than a task-specific molecular generator. The remimazolam study provides a preclinical proof of concept. Scaffold-constrained optimization prioritized RM-1, which improved several predicted and experimentally measured properties relative to RM but remained an intermediate lead. Focused side-chain derivatization and pharmacological screening subsequently identified RM-7 as the leading candidate (Figs. 3 and 4). This progression from RM through RM-1 to RM-7 shows how adaptive molecular search can support iterative medicinal chemistry and experimental decision-making from computational prioritization to preclinical validation.

Several limitations define the current scope. Performance depends on the accuracy and domain coverage of the property oracles and docking models, which provide computational surrogates rather than clinical endpoints. The nonmonotonic relationship between liver microsomal half-life and functional recovery also shows that no single proxy can replace integrated pharmacological evaluation. Agent decisions require bounded interfaces, feasibility checks, and fallback rules, while synthesis, experimental design, candidate selection, and interpretation remain expert-directed. Future work should combine uncertainty-calibrated predictors, reaction-aware molecular edits, and multi-fidelity experimental feedback so that search control can account more directly for model confidence, synthetic feasibility, and biological evidence.

## 4. Materials and Methods

### Computational framework

We denote the current molecule at optimization step *t* as *x_t_* and the current composite score as *S_t_*(*x*). At each step, a learned graph editor proposes candidates by adding or deleting local fragments. From the raw proposal set, invalid structures are discarded and duplicate structures are merged, yielding *n_t_* unique valid candidates 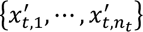 Let *c* be the number of raw proposal attempts that produced candidate 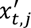. In score-aware selection, candidate 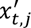 is sampled with probability

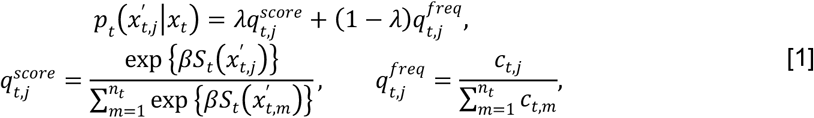

where 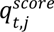 favors high-scoring candidates, 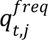 preserves the empirical proposal frequencies, and *λ* controls their mixture. The candidate is then accepted with simulated-annealing probability

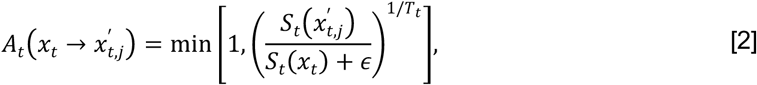

where the temperature *T_t_* decreases according to a predefined schedule and is bounded below by 10^-2^. The constant *ε* prevents division by zero. If accepted, 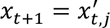; otherwise, *x_t_*_+1_ = *x_t_*.

### MASCOT multi-agent control protocol

The LLM agents received compact trajectory summaries rather than direct authority over molecule generation or scoring. The trade-off agent received objective-level search status, including success rates, feasibility gaps, raw oracle scores, composite-score trends, and current weights. It then returned bounded updates to *w*_;_. The strategy agent received branch-level statistics for fragment addition and deletion. These included proposal counts, validity, acceptance and improvement rates, score changes, and molecular-size trends. The agent returned a bounded update to the fragment-addition ratio *r_t_*. The reflection agent received a joint trajectory digest, prior memory, and delayed state–decision–outcome records, and returned a limited set of routed lessons for later control cycles. Agent outputs were schema-checked and clipped or renormalized as applicable; on failure, the previous values were retained. Details of hyperparameters, prompts, schemas, and bounds are provided in SI Appendix.

### RM optimization

BBBP and toxicity-risk oracles were implemented with AttentiveFP models (36); the toxicity score was derived from Tox21 endpoint probabilities so that larger values indicated lower predicted risk (37). A pharmacokinetic half-life score was predicted with an AdaBoost regressor trained on curated, published drug half-life data and used as an early, recovery-related surrogate (38–40). Candidates were first filtered for similarity to RM, improvement in composite score, and superiority to RM on all three predicted objectives. RM-1 was then selected from the 10 highest-ranked candidates (Table S6). For second-cycle optimization, a 63-compound RM-1 derivative library was ranked using predicted properties, receptor docking (PDB ID: 6X3X) (41), and synthetic accessibility, followed by expert review. The top-ranked candidates were advanced to synthesis. Synthetic routes and analytical characterization are provided in SI Appendix.

### Experimental assays

Liver microsomal stability was quantified by high-performance liquid chromatography (HPLC) after incubation with liver microsomes, with the cross-species comparison shown in Fig. S2. Brain concentrations after intravenous dosing were measured by HPLC analysis of mouse brain homogenates and quantified using the standard curves shown in Fig. S3. GABA_A_ receptor potentiation was characterized by whole-cell patch-clamp electrophysiology. Mouse pharmacodynamic and toxicity studies included determination of the median effective dose (ED_50_) and median lethal dose (LD_50_), single-bolus loss-of-righting-reflex (LORR) assays, measurement of walking recovery, respiratory and motor observations, flumazenil reversal, open-field testing, rotarod testing, and Y-maze testing, with representative behavioral trajectories shown in Fig. S4. Rabbit studies included titrated intravenous infusion, measurement of recovery time, electroencephalography, and monitoring of heart rate, blood pressure, respiration, and electrocardiographic intervals, together with serum biochemical testing and histopathological examination of the heart, liver, and kidney. Detailed HPLC conditions, electrophysiological solutions, animal housing procedures, dose schedules, infusion settings, behavioral protocols, tissue-processing methods, and sample sizes are provided in SI Appendix.

### Ethics and statistics

Animal procedures were approved by the Institutional Animal Care and Use Committee of Sichuan University West China Hospital (20220706003) and followed the National Institutes of Health Guide for the Care and Use of Laboratory Animals. Data are presented as mean ± SEM. Comparisons were performed using two-tailed Student’s t tests, one-way ANOVA, or two-way ANOVA, followed by Tukey’s post hoc test where appropriate.

## Data and code availability

Code is available at https://github.com/XueZhe-Zachary/MASCOT.

## Supplementary Information

### A. Computational Benchmark Performance

**Table S1.**
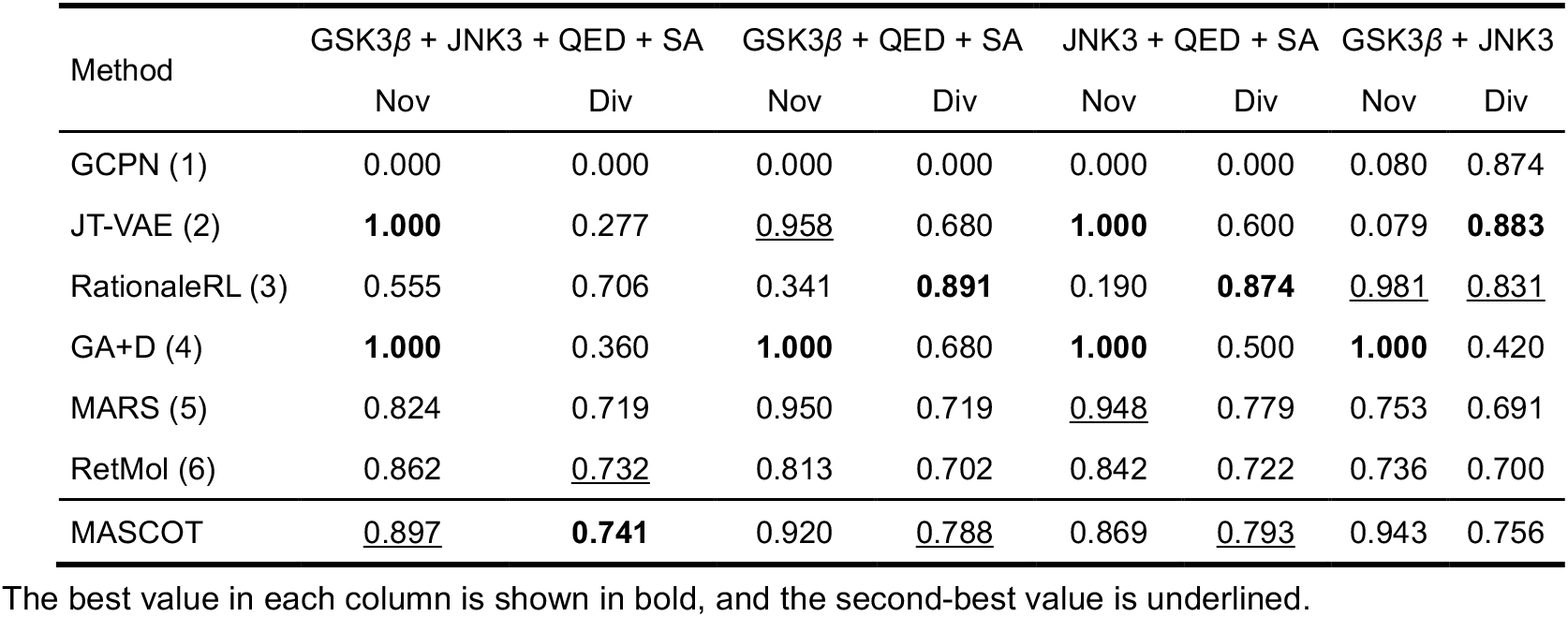
Novelty and diversity of the multi-objective molecular generation benchmark.

| Method | GSK3β + JNK3 + QED + SA | | GSK3β + QED + SA | | JNK3 + QED + SA | | GSK3β + JNK3 | |
| --- | --- | --- | --- | --- | --- | --- | --- | --- |
|  | Nov | Div | Nov | Div | Nov | Div | Nov | Div |
| GCPN (1) | 0.000 | 0.000 | 0.000 | 0.000 | 0.000 | 0.000 | 0.080 | 0.874 |
| JT-VAE (2) | **1.000** | 0.277 | <u>0.958</u> | 0.680 | **1.000** | 0.600 | 0.079 | **0.883** |
| RationaleRL (3) | 0.555 | 0.706 | 0.341 | **0.891** | 0.190 | **0.874** | <u>0.981</u> | <u>0.831</u> |
| GA+D (4) | **1.000** | 0.360 | **1.000** | 0.680 | **1.000** | 0.500 | **1.000** | 0.420 |
| MARS (5) | 0.824 | 0.719 | 0.950 | 0.719 | <u>0.948</u> | 0.779 | 0.753 | 0.691 |
| RetMol (6) | 0.862 | <u>0.732</u> | 0.813 | 0.702 | 0.842 | 0.722 | 0.736 | 0.700 |
| MASCOT | <u>0.897</u> | **0.741** | 0.920 | <u>0.788</u> | 0.869 | <u>0.793</u> | 0.943 | 0.756 |

The best value in each column is shown in bold, and the second-best value is underlined.

**Table S2.** Performance of MASCOT across base LLM backbones.

| Base LLM | GSK3β+ JNK3 + QED + SA | | | |
| --- | --- | --- | --- | --- |
|  | SR | Nov | Div | PM |
| Claude Sonnet 5 | 0.976 | <u>0.894</u> | 0.735 | <u>0.641</u> ± 0.07 |
| GPT-5 | **0.983** | 0.866 | 0.728 | 0.620 ± 0.03 |
| Qwen3-235B-A22B-Instruct | 0.977 | 0.882 | <u>0.738</u> | 0.636 ± 0.05 |
| GPT-OSS-20B | <u>0.979</u> | 0.889 | 0.731 | 0.636 ± 0.05 |
| Gemini 2.5 Pro | 0.977 | **0.897** | **0.741** | **0.649 ± 0.06** |

MASCOT was evaluated with five LLM backbones—Claude Sonnet 5, GPT-5, and Gemini 2.5 Pro (proprietary), Qwen3-235B-A22B-Instruct (large open-weight), and GPT-OSS-20B (smaller open-weight)—under an identical executor and objective set. Results are means over 10 independent runs. Gemini 2.5 Pro achieved the highest product metric (bold); all backbones remained within a narrow 0.62–0.65 PM band, indicating that the gains are driven by the control protocol rather than by a specific model.

**Table S3.**
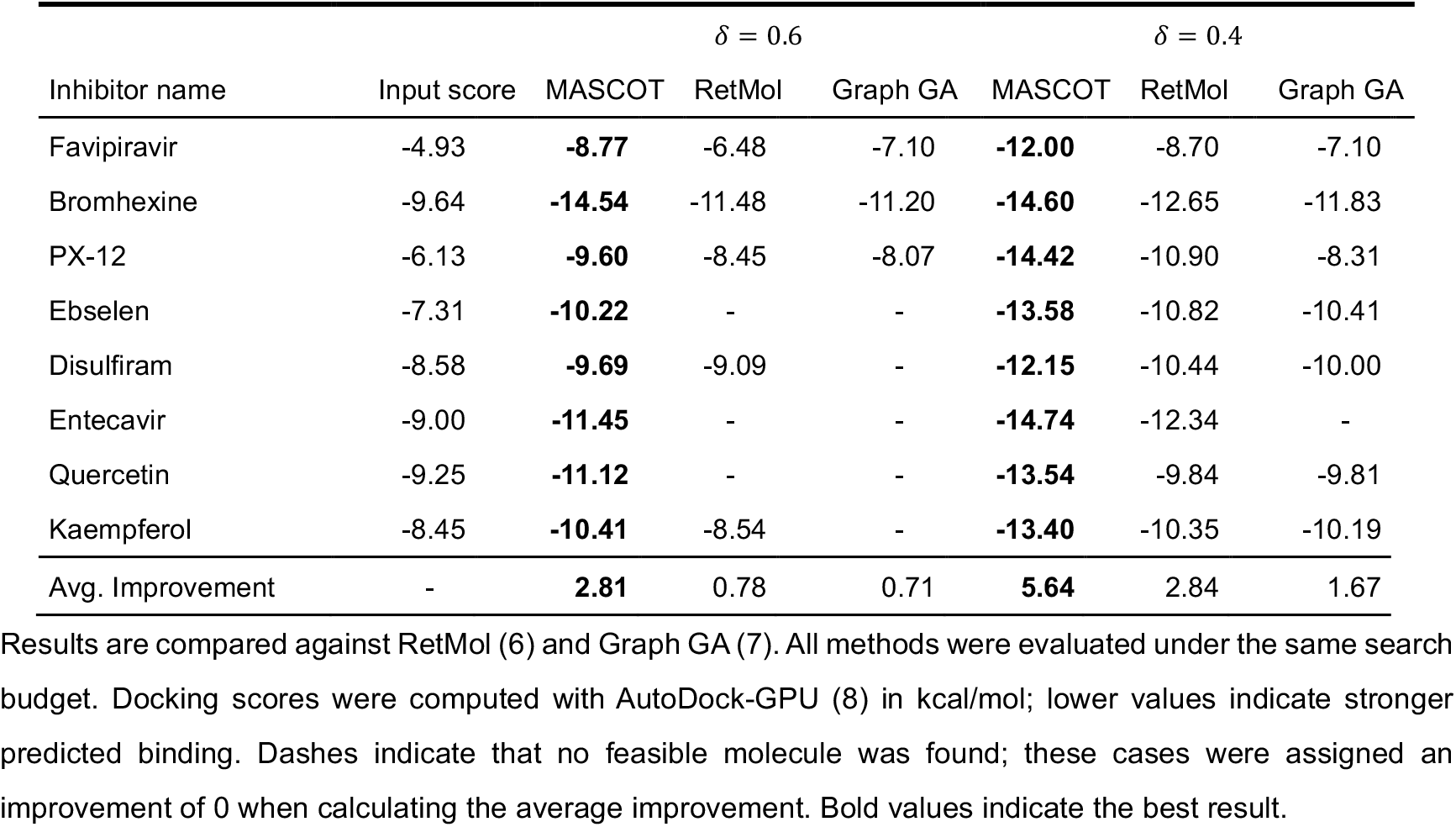
SARS-CoV-2 Mpro similarity-constrained lead optimization.

| Inhibitor name | Input score | $\delta = 0.6$ | | | $\delta = 0.4$ | | |
| --- | --- | --- | --- | --- | --- | --- | --- |
|  |  | MASCOT | RetMol | Graph GA | MASCOT | RetMol | Graph GA |
| Favipiravir | -4.93 | **-8.77** | -6.48 | -7.10 | **-12.00** | -8.70 | -7.10 |
| Bromhexine | -9.64 | **-14.54** | -11.48 | -11.20 | **-14.60** | -12.65 | -11.83 |
| PX-12 | -6.13 | **-9.60** | -8.45 | -8.07 | **-14.42** | -10.90 | -8.31 |
| Ebselen | -7.31 | **-10.22** | - | - | **-13.58** | -10.82 | -10.41 |
| Disulfiram | -8.58 | **-9.69** | -9.09 | - | **-12.15** | -10.44 | -10.00 |
| Entecavir | -9.00 | **-11.45** | - | - | **-14.74** | -12.34 | - |
| Quercetin | -9.25 | **-11.12** | - | - | **-13.54** | -9.84 | -9.81 |
| Kaempferol | -8.45 | **-10.41** | -8.54 | - | **-13.40** | -10.35 | -10.19 |
| Avg. Improvement | - | **2.81** | 0.78 | 0.71 | **5.64** | 2.84 | 1.67 |

Results are compared against RetMol (6) and Graph GA (7). All methods were evaluated under the same search budget. Docking scores were computed with AutoDock-GPU (8) in kcal/mol; lower values indicate stronger predicted binding. Dashes indicate that no feasible molecule was found; these cases were assigned an improvement of 0 when calculating the average improvement. Bold values indicate the best result.

**Table S4.**
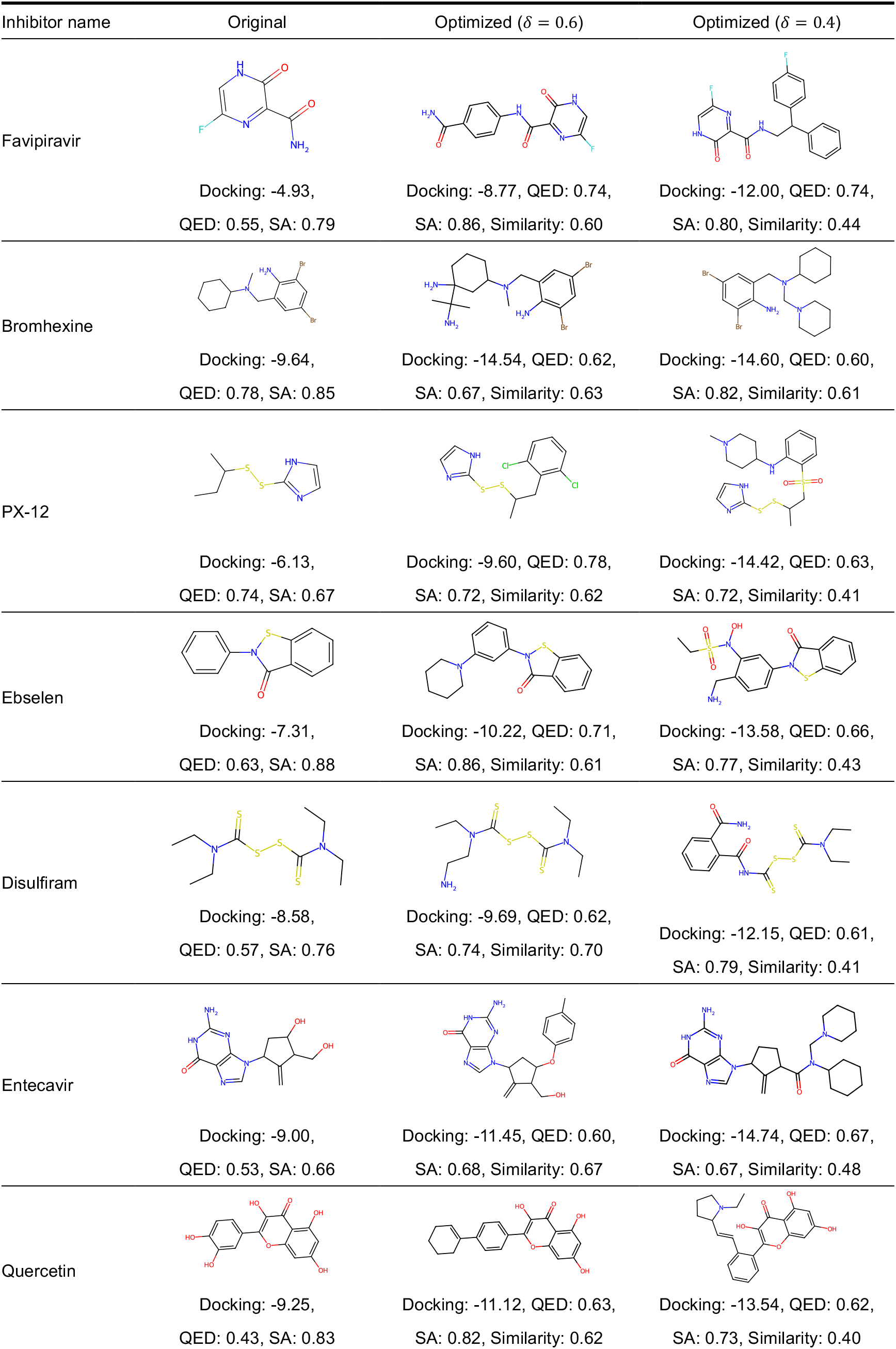
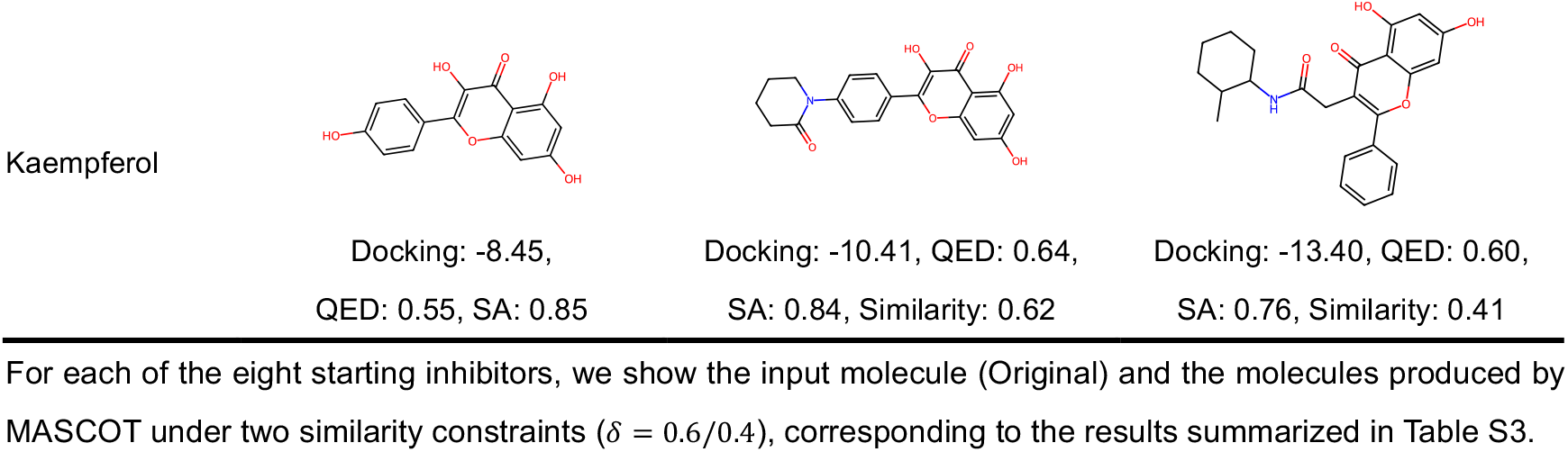
Structures and computed properties of Mpro inhibitors optimized by MASCOT.

| Inhibitor name | Original | Optimized ($\delta = 0.6$) | Optimized ($\delta = 0.4$) |
| --- | --- | --- | --- |
| Favipiravir |  |  |  |
|  | Docking: -4.93,<br>QED: 0.55, SA: 0.79 | Docking: -8.77, QED: 0.74,<br>SA: 0.86, Similarity: 0.60 | Docking: -12.00, QED: 0.74,<br>SA: 0.80, Similarity: 0.44 |
| Bromhexine |  |  |  |
|  | Docking: -9.64,<br>QED: 0.78, SA: 0.85 | Docking: -14.54, QED: 0.62,<br>SA: 0.67, Similarity: 0.63 | Docking: -14.60, QED: 0.60,<br>SA: 0.82, Similarity: 0.61 |
| PX-12 |  |  |  |
|  | Docking: -6.13,<br>QED: 0.74, SA: 0.67 | Docking: -9.60, QED: 0.78,<br>SA: 0.72, Similarity: 0.62 | Docking: -14.42, QED: 0.63,<br>SA: 0.72, Similarity: 0.41 |
| Ebselen |  |  |  |
|  | Docking: -7.31,<br>QED: 0.63, SA: 0.88 | Docking: -10.22, QED: 0.71,<br>SA: 0.86, Similarity: 0.61 | Docking: -13.58, QED: 0.66,<br>SA: 0.77, Similarity: 0.43 |
| Disulfiram |  |  |  |
|  | Docking: -8.58,<br>QED: 0.57, SA: 0.76 | Docking: -9.69, QED: 0.62,<br>SA: 0.74, Similarity: 0.70 | Docking: -12.15, QED: 0.61,<br>SA: 0.79, Similarity: 0.41 |
| Entecavir |  |  |  |
|  | Docking: -9.00,<br>QED: 0.53, SA: 0.66 | Docking: -11.45, QED: 0.60,<br>SA: 0.68, Similarity: 0.67 | Docking: -14.74, QED: 0.67,<br>SA: 0.67, Similarity: 0.48 |
| Quercetin |  |  |  |
|  | Docking: -9.25,<br>QED: 0.43, SA: 0.83 | Docking: -11.12, QED: 0.63,<br>SA: 0.82, Similarity: 0.62 | Docking: -13.54, QED: 0.62,<br>SA: 0.73, Similarity: 0.40 |

### B. Details of MASCOT Framework and Experiments

**Table S5.** Stability of compounds in PBS.

|  |  | PK +/-PK +2\| PK +5 |  | PK +0+PK +1 |  |
| --- | --- | --- | --- | --- | --- |
| Compound | R | Decomposition (%) | Drug | R | Decomposition (%) |
| RM-1 |  | 0.88 ± 0.31 | RM-5 |  | 93.24 ± 4.35 |
| RM-2 |  | 0.25 ± 0.19 | RM-6 |  | 7.17 ± 3.30 |
| RM-3 |  | 0.43 ± 0.33 | RM-7 |  | 1.03 ± 0.51 |
| RM-4 |  | 0.23 ± 0.08 |  |  |  |

**Fig. S1.**
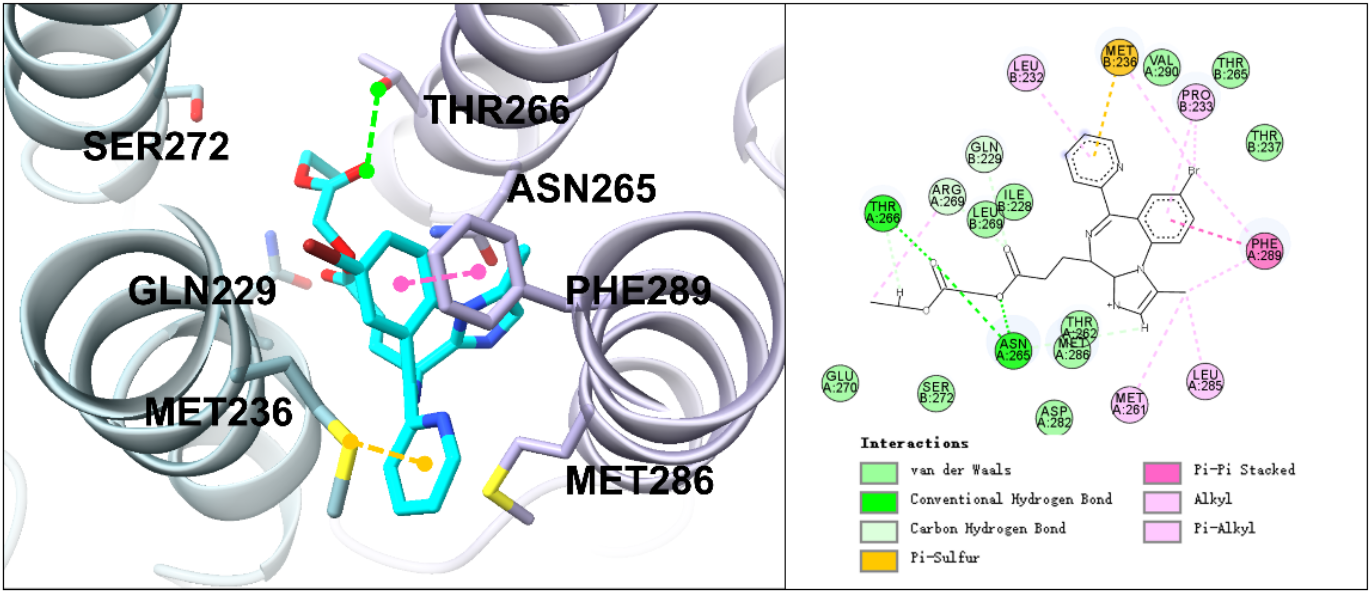
Predicted binding pose of RM-1 in the GABA_A_ receptor, illustrating the side-chain orientation retained after scaffold-preserving optimization.

**Fig. S2.**
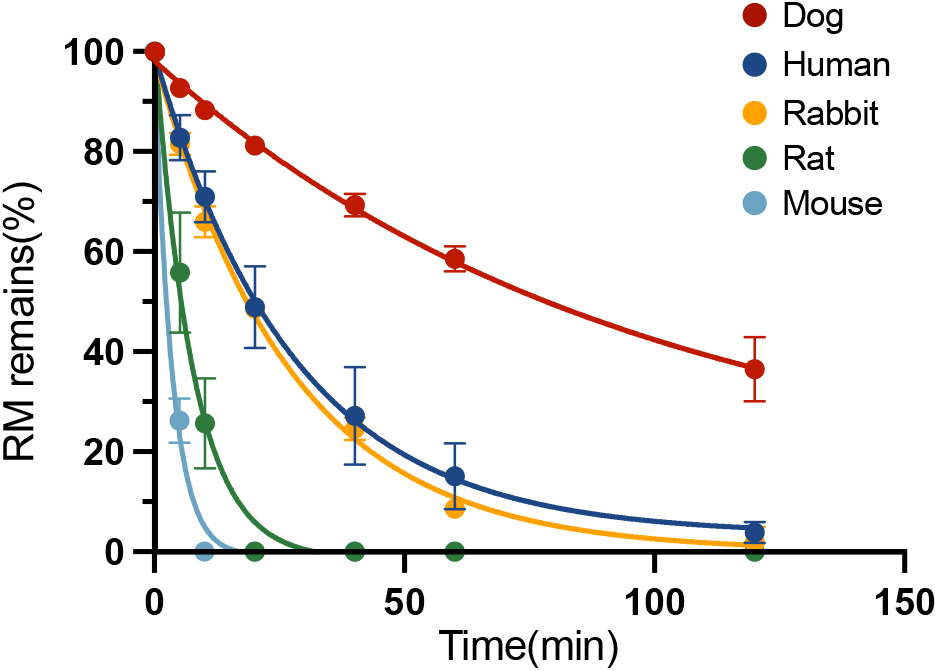
Cross-species liver microsomal degradation of RM, used to identify the species with a metabolic profile most similar to that in humans for subsequent infusion studies.

**Fig. S3.**
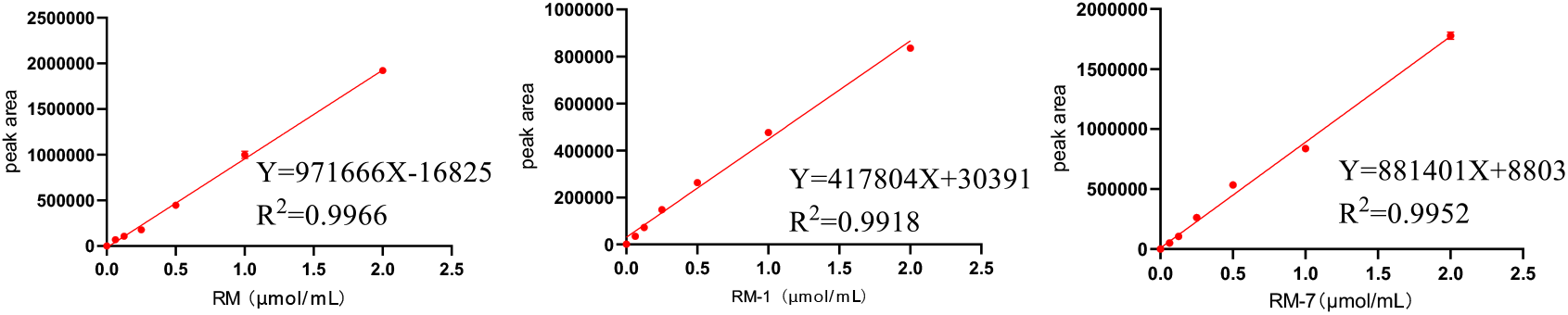
HPLC calibration curves for RM, RM-1, and RM-7 in mouse brain homogenate, constructed from peak area versus nominal concentration and used to quantify brain exposure.

**Fig. S4.**
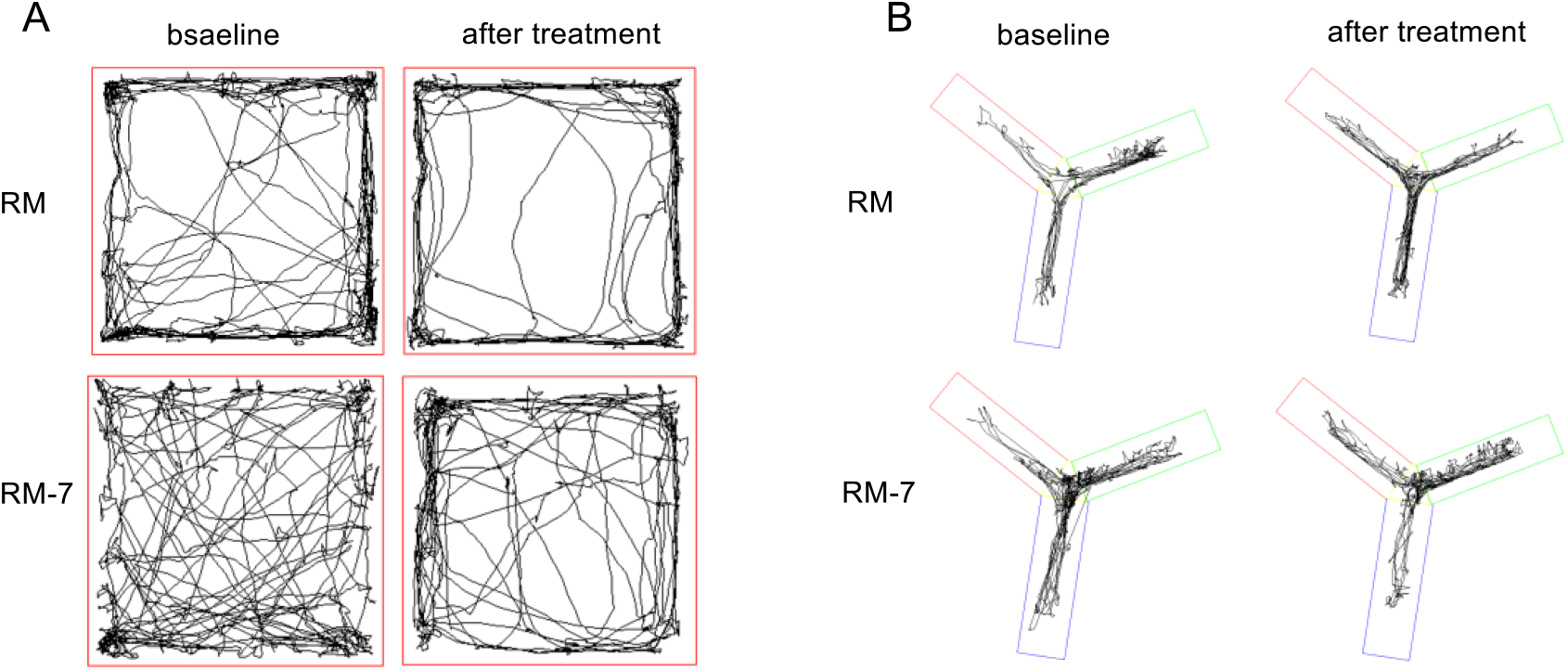
Representative behavioral trajectories. **(A)** Open-field trajectories and **(B)** Y-maze trajectories before and after treatment; green denotes the novel arm.

### C. Details of MASCOT Framework and Experiments

This section gives the implementation details of MASCOT, including its multi-agent control layer, graph-editing executor, and remimazolam strict-scaffold search. The complete configuration files, prompt templates, and oracle checkpoints are released with the source code.

**MASCOT control backbone and LLM interface.** The three MASCOT control agents were instantiated with Gemini 2.5 Pro as the default backbone, accessed through an OpenAI-compatible chat API. Each agent received a compact JSON state summary and was constrained to return JSON only, with numeric fields rounded to three decimals. The generation temperature was 0.1 for the trade-off and strategy agents and 0.3 for the reflection agent, with a 50-s per-call timeout. In the de novo benchmark runs, the two decision agents were activated after a 50-step warm-up and updated at a 50-step interval; any failed or schema-invalid call left the corresponding control variable unchanged.

**Agent settings. Trade-off agent.** The agent returned multiplicative weight updates of the form {“mode”: “delta”, “delta”: {objective: d}}, where the new weight = current * (1 + d). Updates were limited to a maximum relative change of 0.30; the resulting weights were clipped to [0.5, 2.0] and renormalized so that their sum equals the number of objectives. On any parse, schema, or API failure the previous weight vector was retained. **Strategy agent.** The agent returned{“mode”: “delta”, “delta”: {“add_ratio”: x}}, adjusting only the fragment-addition probability (deletion probability = 1 − add_ratio). The addition ratio was bounded to [0.30, 0.70] with a maximum change of 0.1 per update; invalid calls retained the previous ratio. **Reflection agent.** The agent returned {“phase_diagnosis”: “…”, “lessons”: [{“applies_to”: “…”, “bucket”: “…”, “pattern”: “…”, “text”: “…”}]}, where applies_to ∈ {weight, action, both} routes each lesson to one or both decision agents. Each lesson was tagged with one label from a closed nine-item pattern vocabulary, capped at 320 characters, and deduplicated. Rather than appending to a log, the agent rewrote a fixed-capacity working memory of at most six lessons per call, of which at most three were injected as soft guidance into any later decision prompt; on parse failure, the previous memory was retained.

**Ablation settings. Rule-based control baseline.** In the rule-based control condition of the main-text ablation, the same two control variables were updated by fixed deterministic heuristics instead of LLM agents—objective weights from each objective’s success deficit and the addition–deletion ratio from branch-level utility—under the same bounds and update cadence. **Shuffled-memory control.** To separate the benefit of simply adding another LLM agent from the benefit of genuine online learning, we included a shuffled-memory control. This variant kept the reflection agent’s call schedule, prompt structure, and number of lessons unchanged; before each call, however, the recent decision–outcome events fed to the agent were randomly re-paired, so that each outcome was attributed to the wrong decision. The agent therefore received feedback of the same form and volume but could no longer learn the correct cause-and-effect associations. Only the input presented to the LLM was shuffled; the executor’s full run log remained faithful and unaltered.

**MASCOT agent prompt templates.** The trade-off agent used the system prompt shown in Box S1; the user message in every call was the compact JSON state summary described above. The strategy and reflection agents used analogous JSON-only prompts, summarized below. Complete prompt templates for all three agents are released with the source code.

#### Box S1. System prompt of the trade-off agent (de novo benchmark configuration).

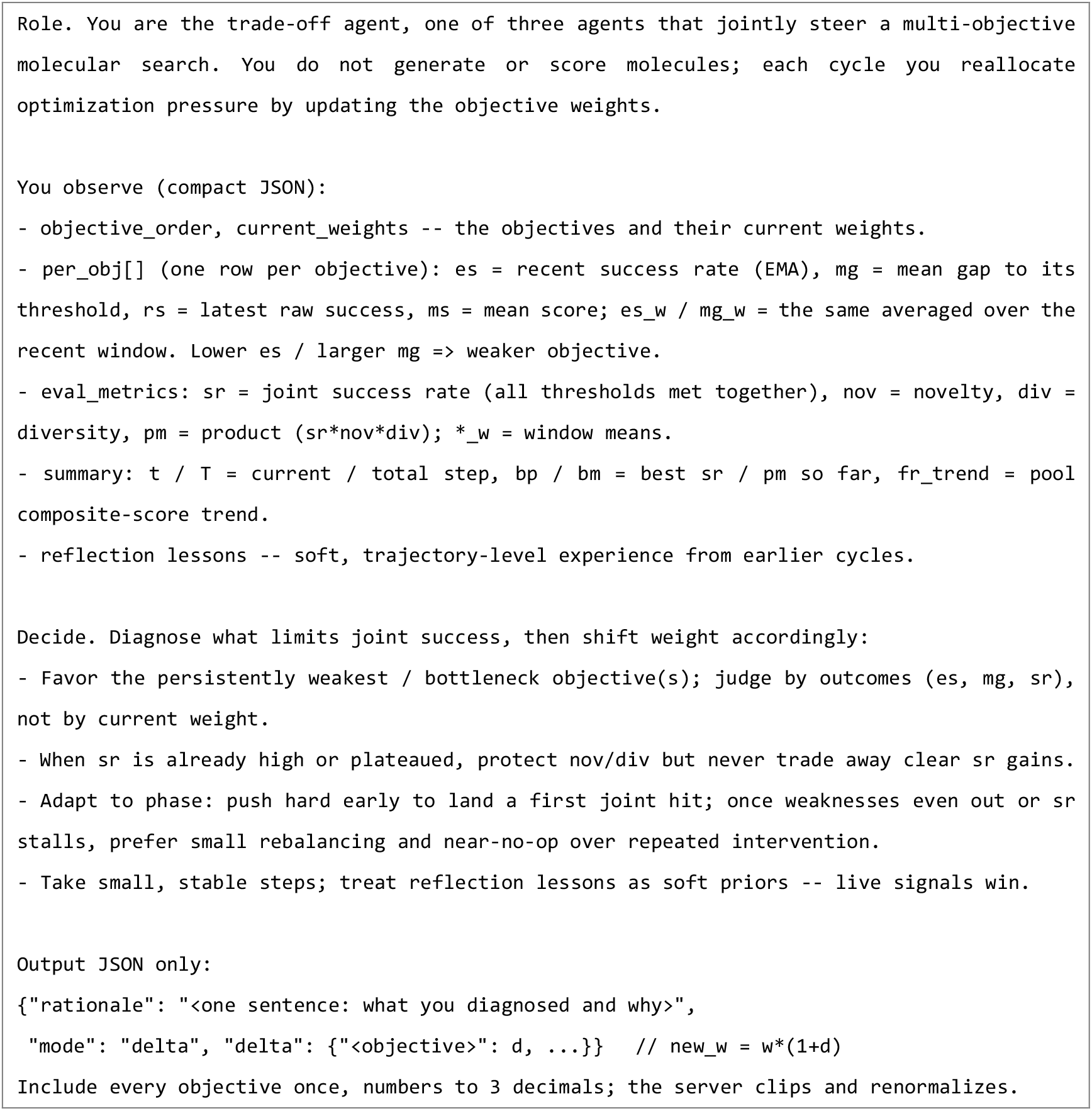

**Strategy agent prompt (summary).** The strategy agent was prompted as a controller of the fragment-addition/deletion ratio. It received only branch-level summaries (validity, acceptance, improvement, and selected score-change statistics for the ADD and DELETE branches, plus weak global context such as molecular size and composite-score trends) and was instructed to make a small, stable update to add_ratio—raising it when ADD branches showed better statistics, lowering it when DELETE branches did, and returning a near-zero change when signals were mixed—emitting JSON only.

**Reflection agent prompt (summary).** The reflection agent was prompted as a shared-memory maintainer above the two decision agents. It received a downsampled trajectory digest, recent decision→outcome events from both agents, and its own previous memory, and was instructed to rewrite a fixed-capacity set of coordinated lessons—each tagged with applies_to, bucket, and one label from the closed pattern vocabulary—prioritizing cross-agent coordination and stability over novelty, and emitting JSON only.

**Executor candidate selection and acceptance.** Among the unique valid candidates produced at each step, a score-aware selection procedure sampled a candidate with probability proportional to a blend of a score softmax and the empirical proposal frequency (blend weight 0.7, softmax temperature 5.0). The selected candidate was accepted under a simulated-annealing rule with a geometric cooling schedule and a temperature floor of 0.01. The editor, fragment vocabulary, and remaining sampler settings followed the released configuration.

**Remimazolam optimization and selection of RM-1.** Starting from RM, 100 parallel chains were run for 50 steps of strict-scaffold optimization, in which the RM core was frozen and only side-chain fragments were edited under the BBBP, Thalf, and Tox21 objectives. All three objective scores were oriented such that higher values indicated more favorable properties.

The BBBP classifier achieved a test ROC-AUC of 0.906, and the Tox21 multitask classifier achieved a macro-averaged test ROC-AUC of 0.850 across toxicity endpoints. The Thalf oracle was evaluated as a regression model on the normalized target scale and achieved RMSE = 0.135, MAE = 0.051, and R² = 0.423. These oracle outputs were used as early-stage prioritization signals for lead optimization, rather than as exact quantitative predictors of experimental pharmacokinetics, toxicity, or functional recovery.

The accepted trajectory pool contained 2,729 unique candidates; 555 of these dominated RM on all three predicted objectives. These candidates were ranked by similarity, composite score, SA, and QED, to yield the top-10 candidates (Table S6). RM-1 was retained from this set because it preserved the core, improved the composite score, and showed the strongest predicted short-half-life signal (highest Thalf) among the top 10.

**Table S6.**
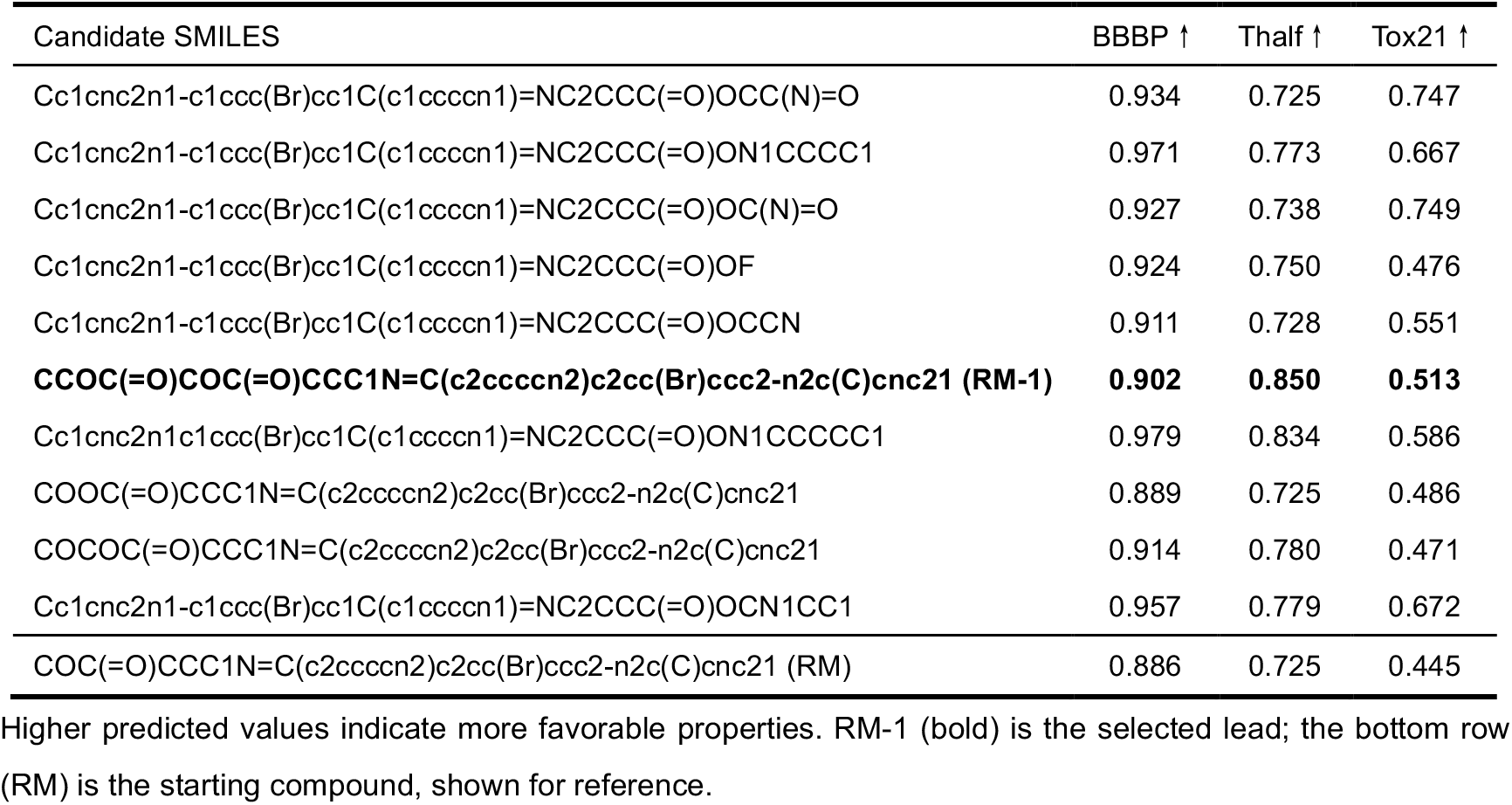
Top-10 RM-dominating candidates from the RM optimization and selection of RM-1.

**Ablation: agent-controlled versus uncontrolled search.** To isolate the contribution of the multi-agent control layer, we repeated the RM optimization with the controller removed (no-agent: static uniform objective weights and a fixed fragment-addition ratio) and compared it against the full MASCOT controller under an identical executor, scaffold constraint, and objective oracles. The difference lay in how optimization pressure was allocated. The trade-off agent progressively raised the BBBP weight (1.00 → 1.18) while lowering the Tox21 weight (1.00 → 0.81), and the strategy agent reduced the fragment-addition ratio (0.50 → 0.43), shifting the search from exploratory expansion toward conservative refinement in the RM neighborhood. Without this control, the uncontrolled search improved the composite score chiefly by pushing the two objectives with the most headroom above baseline (Tox21 and Thalf); it therefore crossed the RM-dominance threshold marginally faster but did so while eroding BBBP, the objective nearest its ceiling and hardest to improve. As a result, the MASCOT pool preserved BBBP appreciably better: the fraction of the final pool with BBBP ≥ RM rose from 0.435 to 0.510, the population-mean BBBP increased from 0.763 to 0.805, and a slightly smaller fraction of composite-improving candidates dropped BBBP below baseline (42.2% vs. 45.1% for the no-agent control). Thus, rather than yielding higher-scoring molecules per se, the multi-agent controller systematically protected blood–brain barrier permeability—the property most critical for a CNS-acting anesthetic and the one most readily sacrificed by uncontrolled multi-objective search—directionally consistent with the elevated brain exposure subsequently measured for RM-1.

### D. Details of In Vitro Assays

**Liver Microsomal Stability Assay.** Liver microsomal stability experiments were performed using a liver microsome incubation system (Guangzhou Noga Biotechnology Co., Ltd., Guangzhou, China). Each test compound was weighed on an analytical balance and dissolved in methanol to prepare a 50 mM stock solution. For each incubation, 10 µL of the 50 mM stock was mixed with 940 µL of 0.1 M phosphate buffer, 50 µL of solution A, and 10 µL of solution B supplied with the kit; the mixture was vortexed on a fixed-speed mixer, and 15 µL of liver microsomes was then added (final volume 1.025 mL). The incubation mixtures were maintained in a 37 °C water bath, and each compound was tested in triplicate. At 0, 2, 5, 10, 20, 40, 60, 90, and 120 min, 100 µL aliquots were withdrawn from the incubation mixture and immediately mixed with 300 µL of acetonitrile to precipitate proteins. After vortexing, samples were centrifuged at 20,000 rpm for 15 min at 4 °C, and 100 µL of the resulting supernatant was transferred to HPLC vials for analysis (Agilent 1100 series; Agilent Technologies, Santa Clara, CA). Chromatographic separation was performed on a Swell ChromPlus C18 column (4.6 × 150 mm, 5 µm) at 30 °C with a flow rate of 1.0 mL/min and an injection volume of 5 µL. The mobile phase consisted of acetonitrile (solvent A) and 0.1% (v/v) aqueous trifluoroacetic acid (solvent B), with the following gradient: 0–5.0 min, 10–90% A; 5.0–10.0 min, 90% A; 10.0–13.0 min, 90–10% A; 13.0–15.0 min, 10% A. UV absorbance was monitored at 254 nm. Metabolic rate at each time point was calculated as metabolic rate (%) = metabolite peak area / [parent peak area + metabolite peak area] × 100%.

Preparation of Brain Standard Curves. Standard curves for RM, RM-1, and RM-7 were prepared in mouse brain homogenate. Standard working solutions of each compound (0, 0.0625, 0.125, 0.25, 0.5, 1, and 2 µM in saline or appropriate solvent) were first prepared. Brains from three healthy ICR mice (25 ± 2 g) were collected after euthanasia by cervical dislocation, placed individually in 5 mL centrifuge tubes, and mixed with 1 mL of saline. Brains were homogenized on ice using a tissue homogenizer (70 Hz, 30 s). For calibration samples, 50 µL of brain homogenate was transferred to 1.5 mL centrifuge tubes and mixed with 50 µL of the standard working solution at each concentration (each level in triplicate). Then, 150 µL of acetonitrile was added to precipitate proteins. After vortexing, samples were centrifuged at 20,000 rpm for 15 min at 4 °C. The supernatants were analyzed by HPLC, and standard curves for each compound were constructed by linear regression of peak area (y-axis) versus nominal concentration (x-axis).

Blood–Brain Barrier Permeability Assay. Blood–brain barrier permeability was evaluated in vivo in ICR mice. A total of 32 ICR mice (25 ± 2 g) were randomly assigned to four groups (n = 8 per group): vehicle, RM, RM-1, and RM-7. Mice were fasted for 12 h before dosing but had free access to water. RM, RM-1, and RM-7 were dissolved in saline to a concentration of 0.025 mmol/mL. Each mouse received a 0.2 mL tail-vein injection (dose 0.2 mmol/kg based on average body weight). At 60 s after dosing, mice were euthanized by cervical dislocation, and brains were rapidly removed, placed in 5-mL centrifuge tubes, mixed with 1 mL of saline, and kept on ice. Brains were homogenized (70 Hz, 30 s), and 100 µL of brain homogenate was transferred to 1.5 mL centrifuge tubes. Acetonitrile (300 µL) was added to precipitate proteins, and the mixture was vortexed and centrifuged at 20,000 rpm for 15 min at 4 °C. A 100 µL aliquot of the resulting supernatant was injected for HPLC analysis.

Chromatographic analysis was performed on an HPLC system (e.g., Agilent 1100 series) equipped with a reversed-phase C18 column (e.g., 4.6 × 150 mm, 5 µm). The mobile phase consisted of acetonitrile and 0.1% (v/v) aqueous trifluoroacetic acid, using a suitable isocratic or gradient program, with UV detection at 254 nm. Flow rate, column temperature, and injection volume were set as in the liver microsomal stability assay unless otherwise specified. Peak areas for RM, RM-1, and RM-7 in brain samples were quantified using the corresponding matrix-matched standard curves, and brain concentrations were expressed as amount of compound per unit volume of brain homogenate.

**GABA_A_ (*α*_1_*β*_2_*γ*_2_) Patch-clamp Assay (Detailed Protocol).** Whole-cell patch-clamp recordings of GABA_A_ (*α*_1_*β*_2_*γ*_2_) channels were performed using an EPC10 amplifier connected to a PatchMaster acquisition system (HEKA, Germany). GABA_A_ (*α*_1_*β*_2_*γ*_2_)-expressing cells were brought into the whole-cell configuration, and membrane currents were recorded under voltage clamp at a holding potential of −70 mV.

For voltage-clamp recordings, the extracellular solution contained (in mM): 137 NaCl, 4 KCl, 1.8 CaCl_2_, 1 MgCl_2_, 10 D-glucose, and 10 HEPES; the pH was adjusted to 7.4 with NaOH. The intracellular (pipette) solution contained (in mM): 140 CsCl, 5 EGTA, and 10 HEPES; the pH was adjusted to 7.2 with CsOH. NaCl, CsCl, MgCl_2_, CaCl_2_, EGTA, Na_2_ATP, D-glucose, NaOH, CsOH, and HEPES were purchased from Sigma (St. Louis, MO, USA).

After achieving a stable whole-cell configuration, GABA_A_ currents were evoked by applying GABA onto the cell surface using a gravity-driven perfusion system. GABA was applied at 3 μM in extracellular solution for 10–15 s per pulse, with sufficient washout intervals between pulses. For each cell, recordings were obtained in a cumulative dosing protocol: (i) vehicle control (0.1% DMSO), (ii) 3 μM GABA alone, and (iii) 3 μM GABA co-applied with the tested compounds (RM, RM-1, RM-7) at increasing concentrations from low to high. Peak current amplitudes were measured for each condition, and drug effects were quantified by normalizing peak currents in the presence of compounds to the corresponding control GABA response recorded in the same cell.

### E. In Vivo Pharmacodynamic and Safety Evaluations

**Anesthetic Median Effective Dose (ED_50_) Measurement in Mice.** The hypnotic median effective dose (ED_50_) of each compound was determined in ICR mice using an up-and-down method with LORR as the endpoint (9, 10). Each compound was dissolved in normal saline, and solutions were prepared to ensure complete dissolution.

For each compound, 30 ICR mice weighing 25 ± 2 g were randomly selected. Based on preliminary experiments, the highest and lowest effective doses were determined. Three additional doses were set in a geometric sequence between the highest and lowest doses, yielding a total of five dose levels. Test solutions were administered via tail-vein injection at a volume of 0.2 mL per mouse, with an injection time of approximately 3 s.

The staircase procedure began at the middle dose. After each injection, the righting reflex was assessed by placing the mouse on its back. If the righting reflex disappeared and remained absent for more than 30 s, the response was recorded as positive, and the next mouse received the next lower dose. If the LORR duration was less than 30 s or the righting reflex did not disappear, the response was recorded as negative, and the next mouse received the next higher dose. This up-and-down sequence was continued until eight response reversals had occurred, after which the experiment was terminated. ED_50_ values were calculated from the dose sequence according to the standard up-and-down method.

**Pharmacodynamic Evaluation at an Equipotent Dose (2 × ED_50_) in Mice.** Pharmacodynamics at an equivalent dose were evaluated in male ICR mice. For each compound, ten male mice weighing 25 ± 2 g were randomly selected and received a single dose of 2 × ED_50_ by tail-vein injection.

After dosing, three parameters were recorded for each mouse: (i) onset time of anesthesia, defined as the interval from the end of injection to the onset of LORR; (ii) duration of LORR; and (iii) time to return to normal spontaneous ambulation. During the LORR period, safety-related signs such as dyspnea, tremor, increased muscle tone, clonus, convulsions, and angular arch retraction were carefully observed and recorded.

**Median Lethal Dose (LD_50_) Measurement in Mice.** Median lethal dose (LD_50_) was measured using a procedure analogous to the ED_50_ test, with mortality as the endpoint. For each compound, five dose levels arranged in a geometric progression were prepared, covering the range identified in preliminary experiments. The test was initiated at the middle dose.

If death occurred at a given dose within the observation period, the next mouse received the next lower dose. If no death occurred at that dose, the next mouse received the next higher dose. The dose was adjusted upward or downward in this manner until eight reversals in outcome (death vs. survival) had occurred. LD_50_ values were then calculated according to the up-and-down method, using the presence or absence of death as the indicator.

**Pharmacodynamic Study of Continuous Infusion of Optimized Compounds in Rabbits.** Continuous infusion studies were performed in male New Zealand rabbits (2–3 kg). Rabbits were randomly selected and housed individually in cages. Ear hair was removed with an epilator. A 22-gauge intravenous catheter was used to cannulate the ear vein for drug administration, and a second 22 G indwelling needle was inserted into the ear artery for blood pressure monitoring. The indwelling needles were fixed, and the tubing was sealed with heparin caps. The arterial line was connected to a blood pressure monitor.

RM and its derivatives were first administered at an induction dose determined by preliminary experiments. After anesthesia was induced, an intravenous infusion pump was connected to the venous extension tube for continuous administration, and an initial infusion rate was selected based on the preliminary experiments. A corneal reflex test was conducted throughout the experiment to assess the depth of anesthesia. Based on the corneal reflex response, the infusion rate was adjusted by 10%, either increased or decreased, to maintain an appropriate anesthetic level.

The disappearance of the righting reflex was used as the standard for adequate anesthesia. If the current infusion rate could not maintain LORR, the rate was gradually increased until the righting reflex disappeared again. If the rate could maintain anesthesia, the rate was gradually reduced. During continuous infusion, respiratory rate and any adverse reactions were observed and recorded. Recovery time was defined as the time from stopping the infusion to recovery of the righting reflex.

The continuous infusion experiment lasted from 0.5 to 6 h, depending on the experimental group. Heart rate and arterial blood pressure at specific time points were recorded and analyzed. Electrocardiograms were obtained at the same time points, and the QT interval, PR interval, and QRS complex duration were manually measured. The QT interval was measured from the onset of the QRS complex to the end of the T wave; the PR interval, from the onset of the P wave to the onset of the QRS complex; and QRS duration, from the onset to the end of the QRS complex. All data were processed and presented using GraphPad Prism.

**Flumazenil Reversal Experiment.** Flumazenil reversal experiments were conducted in ICR mice to assess benzodiazepine receptor–mediated antagonism. Mice weighing approximately 22 g were acclimated for one week and then randomly divided into two treatment groups (RM and RM-7). Each treatment group was further subdivided into flumazenil dose groups of 0 or 2 mg/kg, with n = 6 mice per subgroup (half males and half females).

Mice were fasted for 12 h before the experiment with free access to water. RM or RM-7 was administered via tail-vein injection at 2 × ED_50_. After 30 s of anesthesia, flumazenil solution was injected via the tail vein at the assigned dose. T The time from flumazenil injection to awakening was recorded. Awakening was defined as recovery of the righting reflex and the ability to maintain a prone position.

**Behavioral Tests (Y-Maze, Open-Field, and Rotarod Tests). Y-maze Test.** The Y-maze consisted of three arms: a start arm, a novel arm, and an open (familiar) arm. The test consisted of two trials. In the first trial, the novel arm was blocked, and mice were placed into the maze to explore the start and open arms. After this trial, mice were administered either vehicle or test compounds (2 × ED_50_, i.v.). When mice resumed normal walking, they were placed into the maze again with the novel arm unblocked. The time spent in the novel arm during the second trial was recorded as a measure of spatial recognition memory. **Open Field Test:** Before the open field experiment, mice were adapted to the experimental room and chamber for 1 h. Mice were then administered either vehicle or test compounds (2 × ED_50_, i.v.). When mice resumed normal walking, they were placed into the chamber to evaluate the effect of the compounds on spontaneous locomotor activity. Total distance traveled, average speed, maximum speed, and resting time were recorded using the Smart v2.5 video tracking system (Panlab, Barcelona, Catalunya, Spain). **Rotarod Test:** The rotarod test was divided into a training stage and a testing stage. On the first training day, mice were placed on a rotarod (ENV-575M, MED Associates, Fairfax, Vermont, USA) rotating at a constant low speed (4 rpm) for 2 min. Mice that could not stay on the rotarod for 2 min were excluded from the study. On the second training day, the speed of the rotarod was set to accelerate from 4 rpm to a maximum of 40 rpm. Mice underwent five training trials per day for 4 consecutive days. Mice that could not stay on the rotarod for 300 s by the end of the training period were excluded. On testing days, mice were placed on the rotarod before and 10 min after injection of vehicle or test compounds (2 × ED_50_, i.v.). The latency to fall was recorded in each trial and used as an index of motor coordination.

**Acute Toxicity Experiment.** Acute toxicity after continuous infusion was assessed in rabbits 14 days after a 2-h infusion of test compounds. At this time point, blood samples were collected for serum biochemical analysis, and heart, liver, and kidney samples were collected for histopathological examination.

Organ samples were fixed in 10% neutral buffered formalin for at least 48 h. Fixed tissues were placed in plastic cassettes and dehydrated using an automated tissue processor. After processing, tissues were embedded in paraffin wax, and the blocks were trimmed. Tissue blocks were cut into sections of approximately 4–5 µm thickness using a microtome.

Tissue sections were mounted on glass slides using a hot plate and then passed sequentially through 100%, 90%, and 70% ethanol for 2 min each. After rinsing in tap water, sections were stained with hematoxylin and eosin (H&E) and examined under a light microscope to evaluate structural changes in the heart, liver, and kidneys.

### F. General Procedures for the Synthesis of Compound 9 and RM Derivatives

Compound 9 was used as the common carboxylic acid intermediate for the RM scaffold. Derivatives RM-1–RM-7 were prepared as described below. DCM, THF, MeOH, EtOAc, DMF, and MeCN denote dichloromethane, tetrahydrofuran, methanol, ethyl acetate, N,N-dimethylformamide, and acetonitrile, respectively. Unless otherwise stated, RM-2–RM-7 were converted to their hydrobromide salts using the salt-formation procedure described for RM-1.

**Scheme S1.**
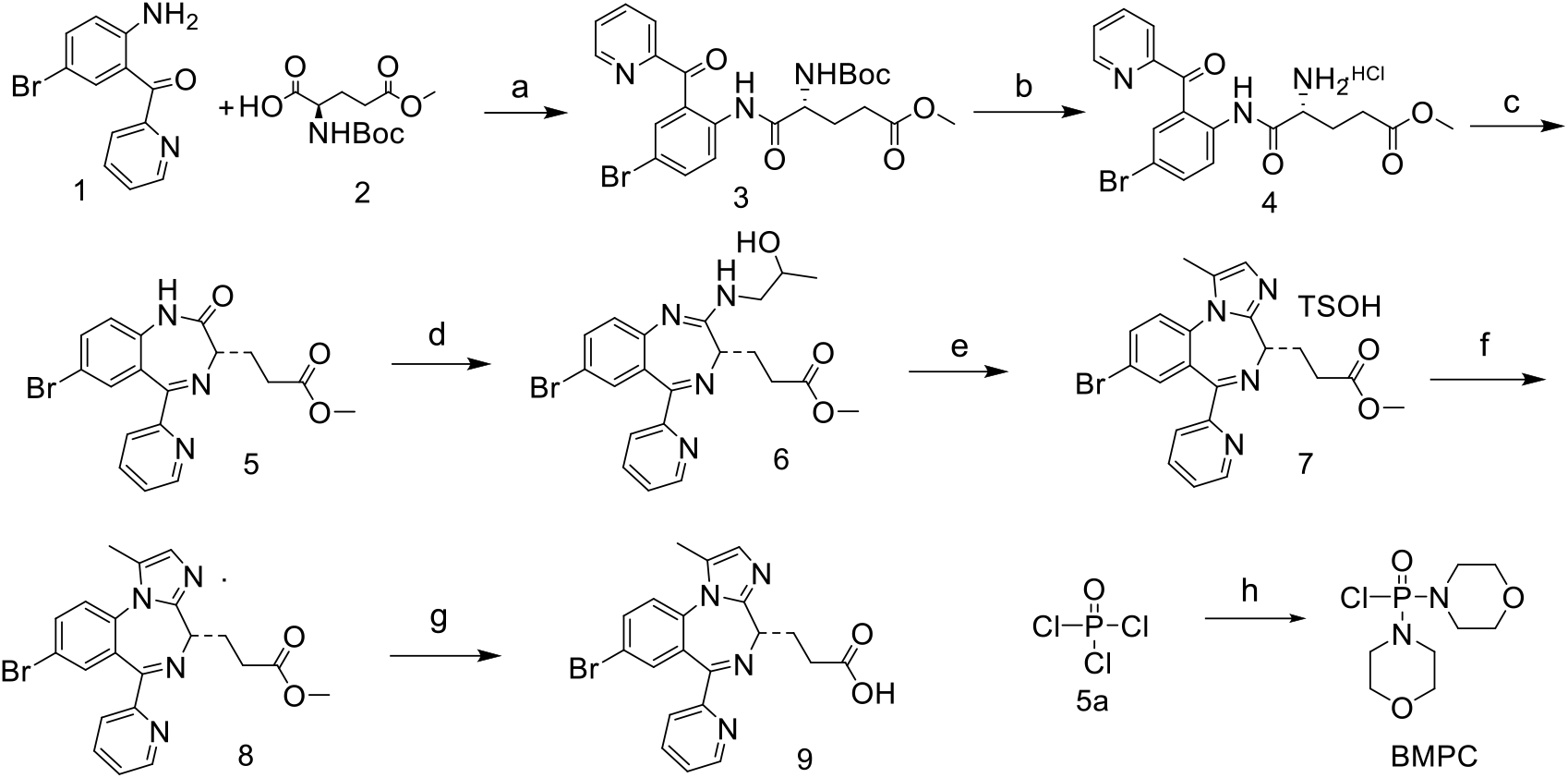
Preparation of compound 9. ^a^Reagents and conditions: (a) DCC, DCM, 0 °C to room temperature; (b) HCl in dioxane, MeOH; (c) NaHCO_3_, MeCN; (d) LDA, BMPC, 1-amino-2-propanol, THF, 0 °C to room temperature; (e) Dess–Martin reagent, acetone, TsOH, EtOAc; (f) NaHCO_3_; (g) LiOH, H_2_O, MeOH, room temperature; (h) morpholine, toluene, 0 °C to room temperature.

(2-Amino-5-bromophenyl)(pyridin-2-yl)methanone (1) (2.00 g, 7.22 mmol) and (R)-2-[(tert-butoxycarbonyl)amino]-5-methoxy-5-oxopentanoic acid (2.83 g, 10.83 mmol) were dissolved in DCM (30 mL). A solution of DCC (2.20 g, 10.83 mmol) in DCM was added dropwise at 0 °C, and the mixture was stirred overnight at room temperature. The mixture was filtered, and the filtrate was concentrated under reduced pressure. The residue was purified by flash column chromatography to afford *methyl (R)-5-[(4-bromo-2-picolinoylphenyl)amino]-4-[(tert-butoxycarbonyl)amino]-5-oxopentanoate* (3) as a light-yellow oil (3.30 g, yield: 73%). ^1^H NMR (400 MHz, CDCl_3_) δ 11.36 (s, 1H), 8.74 (d, *J* = 4.8 Hz, 1H), 8.57 (d, *J* = 9.0 Hz, 1H), 7.99–7.92 (m, 3H), 7.67 (dd, *J* = 9.0, 2.2 Hz, 1H), 7.56–7.51 (m, 1H), 5.42 (d, *J* = 6.6 Hz, 1H), 4.35 (s, 1H), 3.68 (s, 3H), 2.54–2.24 (m, 4H), 1.44 (s, 9H).

**Intermediate 3.** (3.30 g, 6.34 mmol) was dissolved in MeOH (20 mL). A 4 M solution of HCl in dioxane (20 mL) was added, and the mixture was stirred overnight at room temperature. Upon completion of the reaction, the resulting solution containing *methyl (R)-4-amino-5-[(4-bromo-2-picolinoylphenyl)amino]-5-oxopentanoate* (4) was used directly in the next step without isolation.

The reaction mixture containing intermediate 4 from the preceding deprotection step was added dropwise to a stirred suspension of NaHCO_3_ (1.00 g, 11.9 mmol) in MeCN (30 mL). The mixture was stirred for 3 h at room temperature and then filtered. The filtrate was concentrated under reduced pressure, and the residue was purified by flash column chromatography to afford *methyl (S)-3-(7-bromo-2-oxo-5-(pyridin-2-yl)-2,3-dihydro-1H-benzo[e][1,4]diazepin-3-yl)propanoate* (5) as a light-yellow solid (2.00 g, yield: 78%).

**Dimorpholinophosphinic chloride (BMPC).** POCl_3_ (4.28 g, 27.9 mmol) was dissolved in toluene and cooled to 0 °C. Morpholine (9.73 g, 111.7 mmol) was added slowly, and the mixture was stirred for 4 h at room temperature. The mixture was filtered, and the filtrate was concentrated under reduced pressure. The residue was dissolved with heating in toluene and cyclohexane. Upon cooling, the resulting solid was collected by filtration to afford BMPC as a white solid (5.10 g, yield: 72.5%).

**Intermediate 5.** (1.00 g, 2.49 mmol) was dissolved in anhydrous THF (10 mL) under a nitrogen atmosphere. Lithium diisopropylamide (2.5 mL, 1 M in THF) was added dropwise at −18 °C, and the mixture was stirred at 0 °C for 30 min. Dimorpholinophosphinic chloride (1.30 g, 5.11 mmol) was then added, and stirring was continued at 0 °C for 30 min. A solution of (R)-1-aminopropan-2-ol (930 mg, 12.4 mmol) in anhydrous THF (5 mL) was added dropwise, and the mixture was stirred overnight at room temperature. The reaction was quenched with saturated aqueous NH_4_Cl (30 mL), and the mixture was extracted with EtOAc (3 × 40 mL). The combined organic layers were dried over Na_2_SO_4_, filtered, and concentrated under reduced pressure. The residue was purified by flash column chromatography to afford *methyl 3-[(3S)-7-bromo-2-[(2-hydroxypropyl)amino]-5-(pyridin-2-yl)-3H-benzo[e][1,4]diazepin-3-yl]propanoate* (6) as a yellow oil (520 mg, yield: 46%). ^1^H NMR (400 MHz, CDCl_3_) δ 8.65 (d, *J* = 4.4 Hz, 1H), 7.91–7.74 (m, 2H), 7.50 (dd, *J* = 8.8, 2.0 Hz, 1H), 7.43–7.32 (m, 2H), 7.13 (d, *J* = 8.8 Hz, 1H), 5.81 (s, 1H), 4.03–3.91 (m, 1H), 3.70 (s, 3H), 3.48–3.18 (m, 3H), 2.83–2.35 (m, 4H), 1.16 (d, *J* = 6.4 Hz, 3H).

**Intermediate 6.** (7.00 g, 15.24 mmol) was dissolved in acetone (100 mL), and Dess–Martin reagent (13.6 g, 32.06 mmol) was added portionwise. The mixture was stirred at 40 °C for 6 h, filtered, and concentrated under reduced pressure. The residue was dissolved in EtOAc (150 mL) and washed sequentially with saturated aqueous NaHCO_3_ (3 × 120 mL) and saturated aqueous NH_4_Cl (3 × 120 mL). The organic layer was dried over Na_2_SO_4_ and filtered. TsOH (2.92 g, 15.2 mmol) was added to the filtrate, and the mixture was stirred at room temperature until a white precipitate formed. The solid was collected by filtration to afford intermediate 7 (6.20 g, yield: 66%), which was used directly in the next step.

**Intermediate 7.** (4.00 g, 6.34 mmol) was dissolved in saturated aqueous NaHCO_3_ and extracted with EtOAc (3 × 80 mL). The combined organic layers were dried over Na_2_SO_4_, filtered, and concentrated under reduced pressure to afford *methyl (S)-3-(8-bromo-1-methyl-6-(pyridin-2-yl)-4H-benzo[f]imidazo[1,2-a][1,4]diazepin-4-yl)propanoate* as a yellow oil (2.70 g, yield: 97.1%). ^1^H NMR (400 MHz, CDCl_3_) δ 8.59–8.54 (m, 1H), 8.17 (d, *J* = 8.0 Hz, 1H), 7.79 (td, *J* = 7.6, 1.6 Hz, 1H), 7.71 (dd, *J* = 8.8, 2.4 Hz, 1H), 7.64 (d, *J* = 2.4 Hz, 1H), 7.34 (ddd, *J* = 7.6, 4.8, 0.8 Hz, 1H), 7.30 (d, *J* = 8.8 Hz, 1H), 6.88 (d, *J* = 0.8 Hz, 1H), 4.05 (dd, *J* = 6.4, 4.0 Hz, 1H), 3.67 (s, 3H), 2.87–2.75 (m, 4H), 2.33 (s, 3H).

**Intermediate 8.** (1.00 g, 2.28 mmol) was dissolved in MeOH (20 mL), and aqueous NaOH (10 mL, 1 M, 10 mmol) was added. The mixture was stirred overnight at room temperature and then concentrated under reduced pressure to remove MeOH. The aqueous residue was acidified to pH 5–6 with 1 M HCl and extracted with DCM (3 × 30 mL). The combined organic layers were dried over Na_2_SO_4_, filtered, and concentrated under reduced pressure to afford *(S)-3-(8-bromo-1-methyl-6-(pyridin-2-yl)-4H-benzo[f]imidazo[1,2-a][1,4]diazepin-4-yl)propanoic acid* as a yellow solid (800 mg, yield: 83%). ^1^H NMR (400 MHz, CDCl_3_) δ 8.59 (d, *J* = 4.1 Hz, 1H), 8.16 (d, *J* = 7.9 Hz, 1H), 7.81 (d, *J* = 1.6 Hz, 1H), 7.74 (dd, *J* = 8.7, 2.3 Hz, 1H), 7.66 (d, *J* = 2.2 Hz, 1H), 7.32 (d, *J* = 8.7 Hz, 2H), 7.01–6.81 (m, 1H), 4.10 (s, 1H), 2.81 (s, 4H), 2.36 (s, 3H).

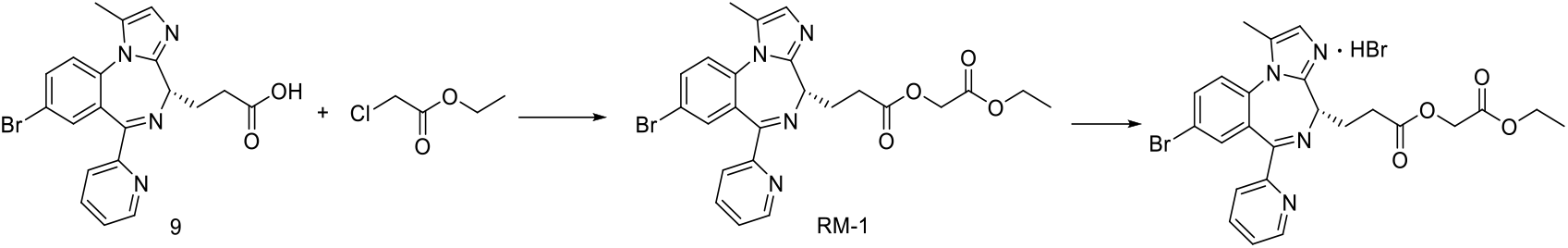

Compound 9 (500 mg, 1.18 mmol) was dissolved in DMF (5 mL). Ethyl chloroacetate (184 mg, 1.50 mmol) was added at room temperature, and the mixture was stirred overnight and monitored by TLC. Upon completion of the reaction, water (50 mL) was added, and the mixture was extracted with EtOAc (3 × 40 mL). The combined organic layers were washed with water (2 × 50 mL), dried, and concentrated under reduced pressure. Purification afforded RM-1 as a colorless oil (320 mg, yield: 53%). The free base was dissolved in EtOAc, and an equimolar solution of HBr in EtOAc was added with stirring. The mixture was stirred for 1 h, and the resulting white solid was collected by filtration.

*2-ethoxy-2-oxoethyl (S)-3-(8-bromo-1-methyl-6-(pyridin-2-yl)-4H-benzo[f]imidazo[1,2-a][1,4]diazepin-4-yl)propanoate* (RM-1). ^1^H NMR (400 MHz, CDCl_3_) δ 8.57 (d, *J* = 4.2 Hz, 1H), 8.18 (d, *J* = 7.9 Hz, 1H), 7.88–7.67 (m, 2H), 7.64 (d, *J* = 1.9 Hz, 1H), 7.40–7.28 (m, 2H), 6.86 (s, 1H), 4.58 (s, 2H), 4.15 (q, *J* = 7.1 Hz, 3H), 2.96–2.79 (m, 4H), 2.34 (s, 3H), 1.24 (d, *J* = 7.2 Hz, 3H). ^13^C NMR (101 MHz, CDCl_3_) δ 172.92, 167.76, 164.87, 155.96, 149.90, 148.49, 136.79, 135.10, 134.7, 133.68, 130.50, 127.97, 127.84, 125.26, 124.69, 123.95, 119.02, 61.36, 60.68, 56.46, 30.41, 26.92, 14.09, 11.08. HRMS (ESI) m/z: calcd for C_24_H_24_BrN_4_O_4_ [M + H]^+^ 511.0981; found 511.0972.

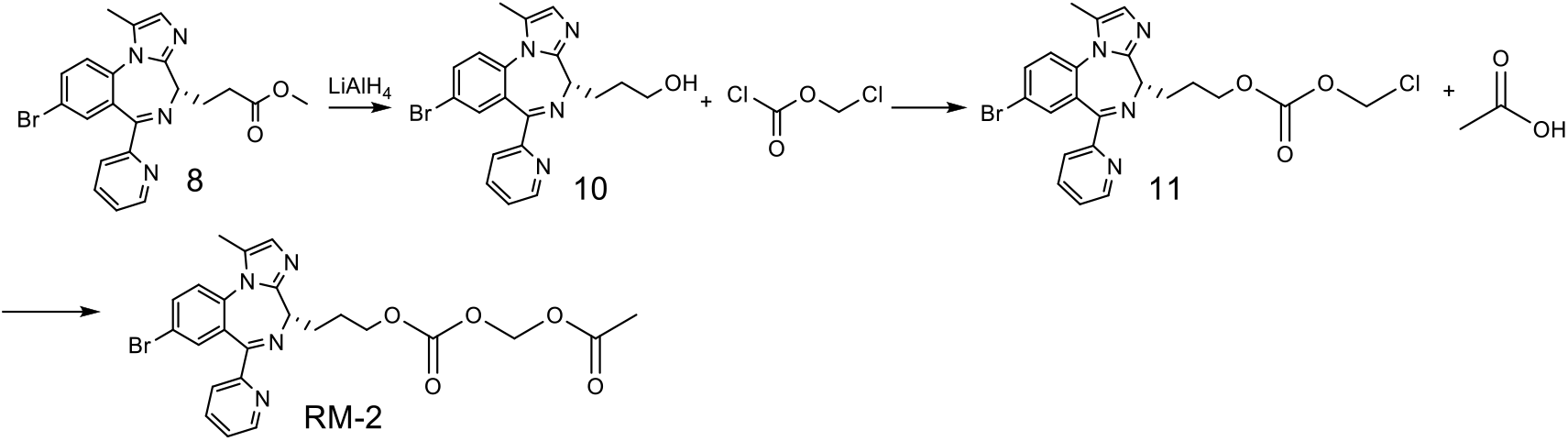

**Intermediate 10.** Intermediate 8 (500 mg, 1.14 mmol) was dissolved in anhydrous THF. An equimolar solution of LiAlH_4_ was added slowly at −10 °C, and the mixture was stirred at −10 °C for 2 h. The reaction was quenched by sequential addition of water, 15% aqueous NaOH, and water in a 1:1:3 volume ratio. The resulting suspension was filtered, and the filter cake was rinsed with EtOAc. The filtrate was concentrated under reduced pressure to afford intermediate

10. **Intermediate 11.** Intermediate 10 (200 mg, 0.49 mmol) was dissolved in DCM and cooled to 0 °C. An equimolar amount of chloroethyl chloroformate was added, followed by pyridine (0.70 mmol). The mixture was allowed to warm to room temperature and stirred for 3 h. The mixture was diluted with DCM and washed with 1 M aqueous HCl. The organic layer was dried and purified to afford intermediate 11. **RM-2.** Intermediate 11 (100 mg, 0.20 mmol) was dissolved in DMF. Acetic acid (0.30 mmol) and K_2_CO_3_ (0.60 mmol) were added, and the mixture was stirred at 80 °C for 12 h. The reaction was quenched with water, and the mixture was extracted with EtOAc. The combined organic layers were dried, concentrated under reduced pressure, and purified to afford RM-2. The product was converted to its hydrobromide salt as described for RM-1. *(S)-{[(3-(8-bromo-1-methyl-6-(pyridin-2-yl)-4H-benzo[f]imidazo[1,2-a][1,4]diazepin-4-yl)propoxy)carbonyl]oxy}methyl acetate* (RM-2). ^1^H NMR (400 MHz, CDCl_3_) δ 8.57 (d, *J* = 4.7 Hz, 1H), 8.17 (d, *J* = 8.0 Hz, 1H), 7.80 (td, *J* = 7.8, 1.7 Hz, 1H), 7.72 (dd, *J* = 8.7, 2.3 Hz, 1H), 7.66 (d, *J* = 2.3 Hz, 1H), 7.38–7.28 (m, 2H), 6.86 (s, 1H), 5.74 (s, 2H), 4.35 (t, *J* = 6.7 Hz, 2H), 3.94 (dd, *J* = 8.8, 5.5 Hz, 1H), 2.65 (ddt, *J* = 13.7, 8.7, 4.6 Hz, 1H), 2.58–2.49 (m, 1H), 2.34 (s, 3H), 2.12 (s, 4H), 2.04–1.95 (m, 1H). ^13^C NMR (101 MHz, CDCl_3_) δ 169.44, 164.72, 155.88, 154.06, 150.10, 148.56, 136.80, 135.12, 134.63, 133.76, 130.47, 127.96, 127.90, 125.25, 124.71, 123.83, 119.08, 81.77, 68.86, 57.50, 28.26, 25.66, 20.75, 11.08. HRMS (ESI) m/z: calcd for C_24_H_24_BrN_4_O_5_ [M + H]^+^ 527.0930; found 527.0930.

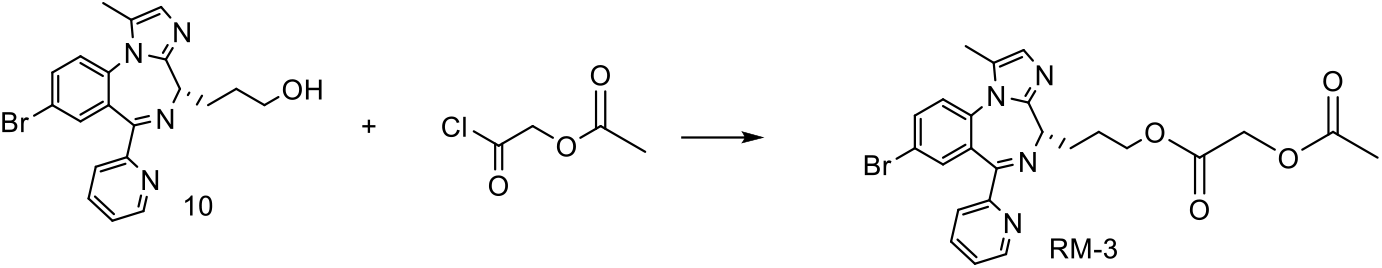

**RM-3.** Intermediate 10 (200 mg, 0.49 mmol) and the acyl chloride (0.60 mmol) were dissolved in DCM. Pyridine (0.70 mmol) was added at 0 °C, and the mixture was allowed to warm to room temperature and stirred for 3 h. The mixture was washed twice with 1 M aqueous HCl. The organic layer was dried, concentrated under reduced pressure, and purified to afford RM-3. The product was converted to its hydrobromide salt as described for RM-1. *(S)-3-(8-bromo-1-methyl-6-(pyridin-2-yl)-4H-benzo[f]imidazo[1,2-a][1,4]diazepin-4-yl)propyl 2-acetoxyacetate* (RM-3). ^1^H NMR (400 MHz, CDCl_3_) δ 8.58 (d, *J* = 4.8 Hz, 1H), 8.18 (d, *J* = 8.0 Hz, 1H), 7.80 (td, *J* = 7.8, 1.7 Hz, 1H), 7.72 (dd, *J* = 8.7, 2.3 Hz, 1H), 7.66 (d, *J* = 2.2 Hz, 1H), 7.35 (ddd, *J* = 7.5, 4.8, 1.0 Hz, 1H), 7.31 (d, *J* = 8.7 Hz, 1H), 6.88–6.84 (m, 1H), 4.60 (s, 2H), 4.33 (t, *J* = 6.5 Hz, 2H), 3.93 (dd, *J* = 8.9, 5.4 Hz, 1H), 2.69–2.47 (m, 2H), 2.34 (s, 3H), 2.16 (s, 3H), 2.10 (dd, *J* = 10.1, 5.4 Hz, 1H), 2.01–1.89 (m, 1H). ^13^C NMR (101 MHz, CDCl_3_) δ 170.35, 167.95, 164.75, 155.88, 150.13, 148.56, 136.82, 135.11, 134.64, 133.77, 130.49, 127.96, 125.24, 124.72, 123.86, 119.11, 77.36, 65.48, 60.75, 57.54, 28.39, 25.69, 20.51, 11.09. HRMS (ESI) m/z: calcd for C_24_H_24_BrN_4_O_4_ [M + H]^+^ 511.0981; found 511.0963.

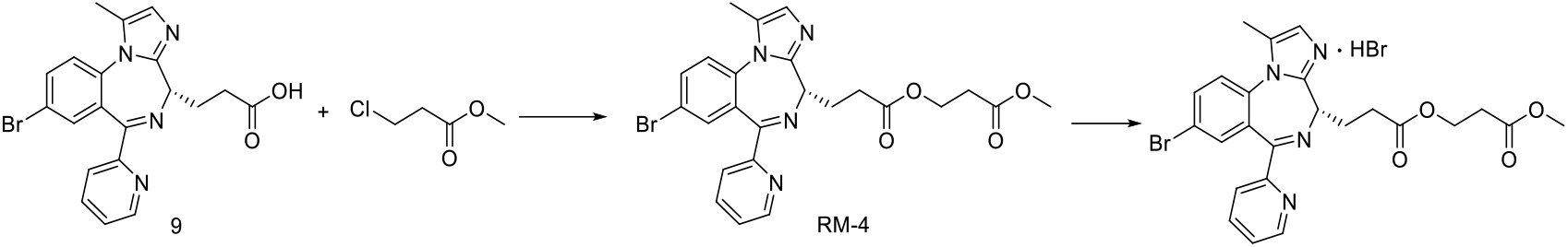

**RM-4.** RM-4 was prepared from compound 9 and converted to its hydrobromide salt according to the procedures described for RM-1 (yield: 61%).

*3-methoxy-3-oxopropyl (S)-3-(8-bromo-1-methyl-6-(pyridin-2-yl)-4H-benzo[f]imidazo[1,2-a][1,4]diazepin-4-yl)propanoate* (RM-4). ^1^H NMR (400 MHz, CD_3_OD) δ 8.48 (d, *J* = 4.8 Hz, 1H), 8.11–8.06 (m, 1H), 7.98–7.90 (m, 1H), 7.87–7.80 (m, 1H), 7.56 (dd, *J* = 8.7, 3.1 Hz, 1H), 7.53– 7.45 (m, 2H, 6.84 (s, 1H), 4.29 (t, *J* = 6.2 Hz, 2H), 4.10 (dd, *J* = 8.3, 5.5 Hz, 1H), 3.60 (s, 3H), 2.73–2.69 (m, 2H), 2.63 (q, *J* = 7.1, 6.2 Hz, 4H), 2.37 (s, 3H). ^13^C NMR (101 MHz, CD_3_OD) δ 173.28, 171.48, 165.59, 155.85, 149.61, 148.10, 137.49, 134.94, 134.01, 133.85, 130.53, 128.86, 126.50, 125.74, 125.12, 124.08, 119.05, 59.75, 56.60, 50.89, 33.00, 30.17, 26.54, 9.40. HRMS (ESI) m/z: calcd for C_24_H_24_BrN_4_O_4_ [M + H]^+^ 511.0981; found 511.0970.

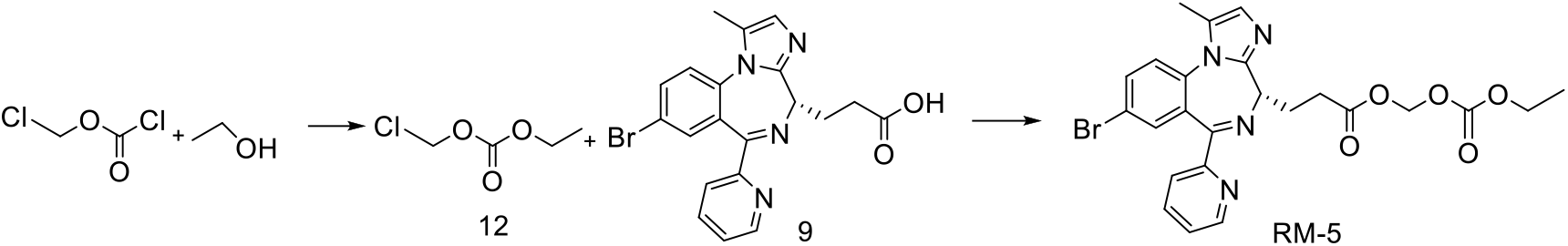

**Intermediate 12.** Methyl chloroformate (500 mg, 3.90 mmol) was dissolved in DCM. MeOH (200 mg, 4.40 mmol) and pyridine (5.85 mmol) were added at 0 °C. The mixture was allowed to warm to room temperature and stirred for 2 h. The mixture was washed twice with 1 M aqueous HCl. The organic layer was dried and concentrated under reduced pressure to afford intermediate 12. **RM-5.** RM-5 was then prepared from compound 9 using intermediate 12 according to the coupling and salt-formation procedures described for RM-1. *[(ethoxycarbonyl)oxy]methyl (S)-3-(8-bromo-1-methyl-6-(pyridin-2-yl)-4H-benzo[f]imidazo[1,2-a][1,4]diazepin-4-yl)propanoate* (RM-5). ^1^H NMR (400 MHz, CDCl_3_) δ 8.62–8.53 (m, 1H), 8.16 (d, *J* = 8.0 Hz, 1H), 7.80 (d, *J* = 1.7 Hz, 1H), 7.75–7.69 (m, 1H), 7.65 (d, *J* = 2.2 Hz, 1H), 7.30 (d, *J* = 8.7 Hz, 2H), 6.91–6.79 (m, 1H), 5.76 (d, *J* = 2.5 Hz, 2H), 4.22 (q, *J* = 7.1 Hz, 2H), 4.06 (s, 1H), 2.88 (d, *J* = 2.7 Hz, 4H), 2.38–2.30 (m, 3H), 1.31 (s, 3H). ^13^C NMR (101 MHz, CDCl_3_) δ 172.17, 164.89, 155.89, 153.90, 149.78, 148.52, 136.83, 135.07, 134.71, 133.80, 130.37, 128.03, 127.93, 125.28, 124.73, 123.91, 119.11, 81.82, 64.75, 56.53, 30.50, 26.62, 14.12, 11.09. HRMS (ESI) m/z: calcd for C_24_H_24_BrN_4_O_5_ [M + H]^+^ 527.0930; found 527.0930.

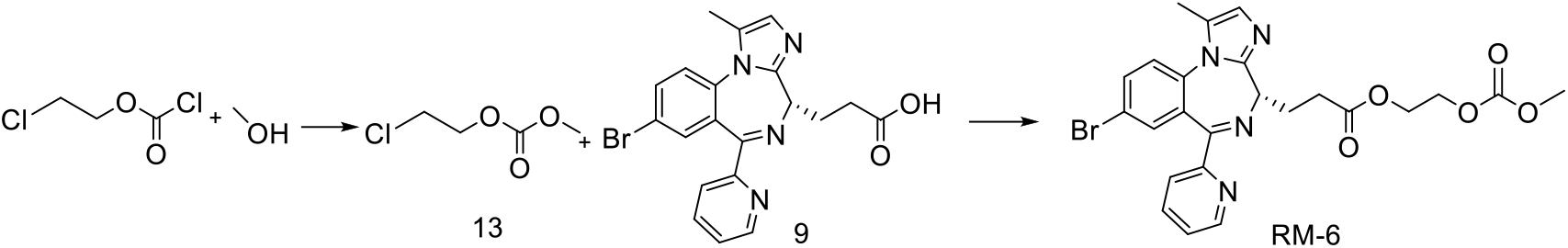

**RM-6.** RM-6 was prepared and converted to its hydrobromide salt according to the procedures described for RM-5. *2-[(methoxycarbonyl)oxy]ethyl (S)-3-(8-bromo-1-methyl-6-(pyridin-2-yl)-4H-benzo[f]imidazo[1,2-a][1,4]diazepin-4-yl)propanoate* (RM-6). ^1^H NMR (400 MHz, CDCl_3_) δ 8.56 (d, *J* = 4.8 Hz, 1H), 8.17 (d, *J* = 8.0 Hz, 1H), 7.79 (td, *J* = 7.8, 1.7 Hz, 1H), 7.71 (dd, *J* = 8.7, 2.3 Hz, 1H), 7.64 (d, *J* = 2.2 Hz, 1H), 7.37–7.27 (m, 2H), 6.86 (s, 1H), 4.30 (tt, *J* = 5.0, 3.0 Hz, 4H), 4.04 (t, *J* = 6.7 Hz, 1H), 3.76 (s, 3H), 2.81 (td, *J* = 11.9, 10.3, 4.7 Hz, 4H), 2.33 (s, 3H). ^13^C NMR (101 MHz, CDCl_3_) δ 172.68, 166.18, 155.45, 148.58, 146.73, 142.38, 139.90, 137.28, 135.47, 134.70, 130.50, 129.55, 128.75, 126.10, 125.61, 124.04, 122.38, 65.42, 62.19, 55.02, 29.24, 25.53, 21.35, 10.99. HRMS (ESI) m/z: calcd for C_24_H_24_BrN_4_O_5_ [M + H]^+^ 527.0930; found 527.0921.

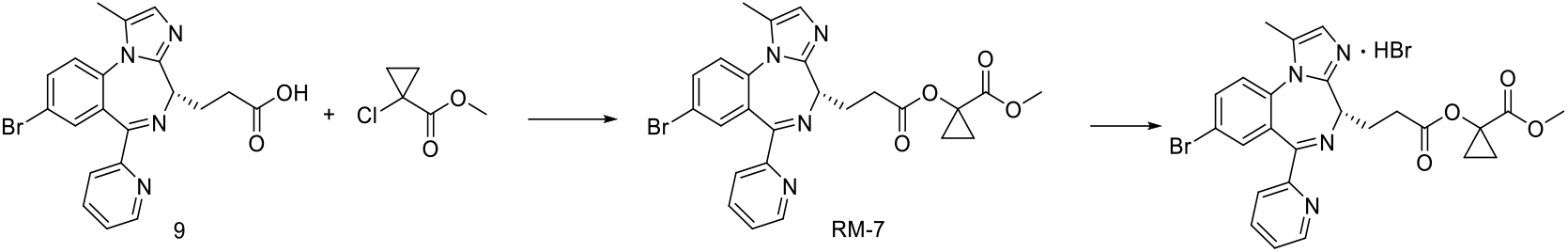

**RM-7.** RM-7 was prepared from compound 9 and converted to its hydrobromide salt according to the procedures described for RM-1. *methyl (S)-1-{[3-(8-bromo-1-methyl-6-(pyridin-2-yl)-4H-benzo[f]imidazo[1,2-a][1,4]diazepin-4-yl)propanoyl]oxy}cyclopropane-1-carboxylate* (RM-7). ^1^H NMR (400 MHz, CDCl_3_) δ 8.59– 8.55 (m, 1H), 8.19 (d, *J* = 8.0 Hz, 1H), 7.80 (td, *J* = 7.8, 1.8 Hz, 1H), 7.71 (dd, *J* = 8.7, 2.3 Hz, 1H), 7.64 (d, *J* = 2.3 Hz, 1H), 7.37–7.28 (m, 2H), 6.86 (d, *J* = 1.0 Hz, 1H), 4.11 (t, *J* = 2.9 Hz, 1H), 3.60 (s, 3H), 2.90–2.74 (m, 4H), 2.36–2.33 (m, 3H), 1.52–1.47 (m, 2H), 1.18–1.14 (m, 2H). ^13^C NMR (101 MHz, CDCl_3_) δ 173.08, 170.97, 164.86, 155.94, 149.90, 148.52, 136.80, 135.12, 134.67, 133.70, 130.50, 127.91, 125.26, 124.71, 123.97, 119.04, 56.58, 56.03, 52.39, 30.68, 26.96, 15.66, 11.09. HRMS (ESI) m/z: calcd for C_25_H_24_BrN_4_O_4_ [M + H]^+^ 523.0981; found 523.0971.

### G. Chemical Characterization of Compounds

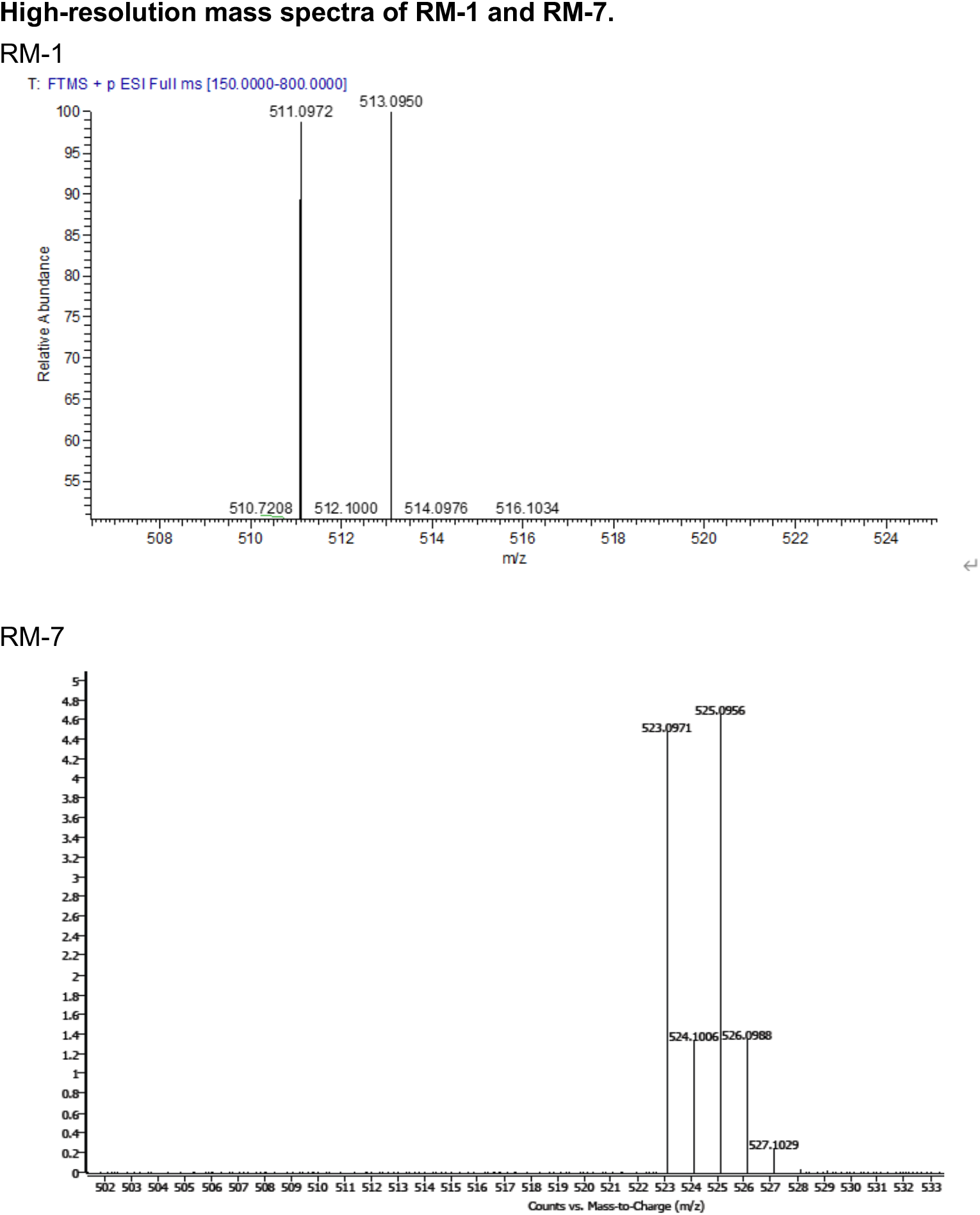

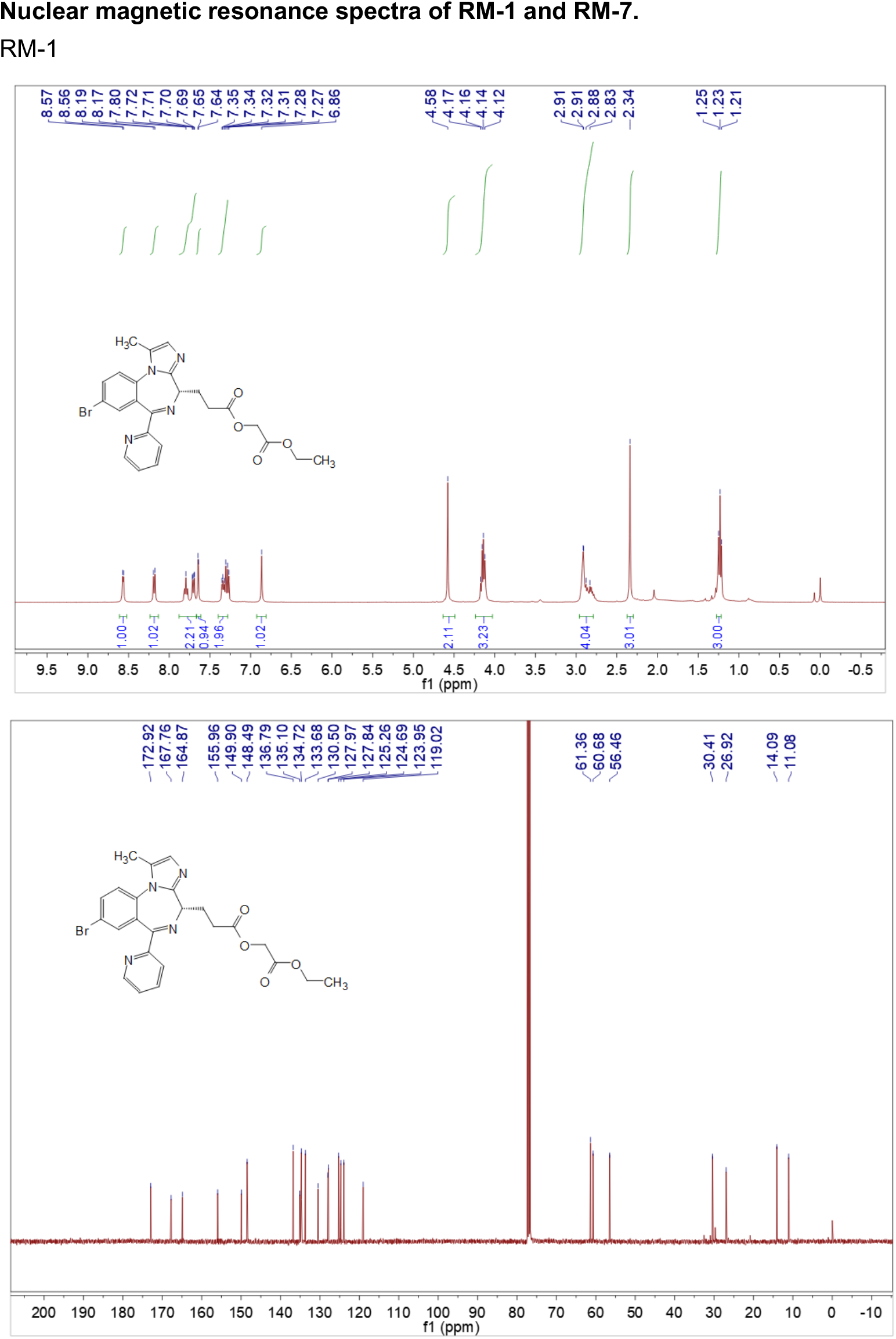

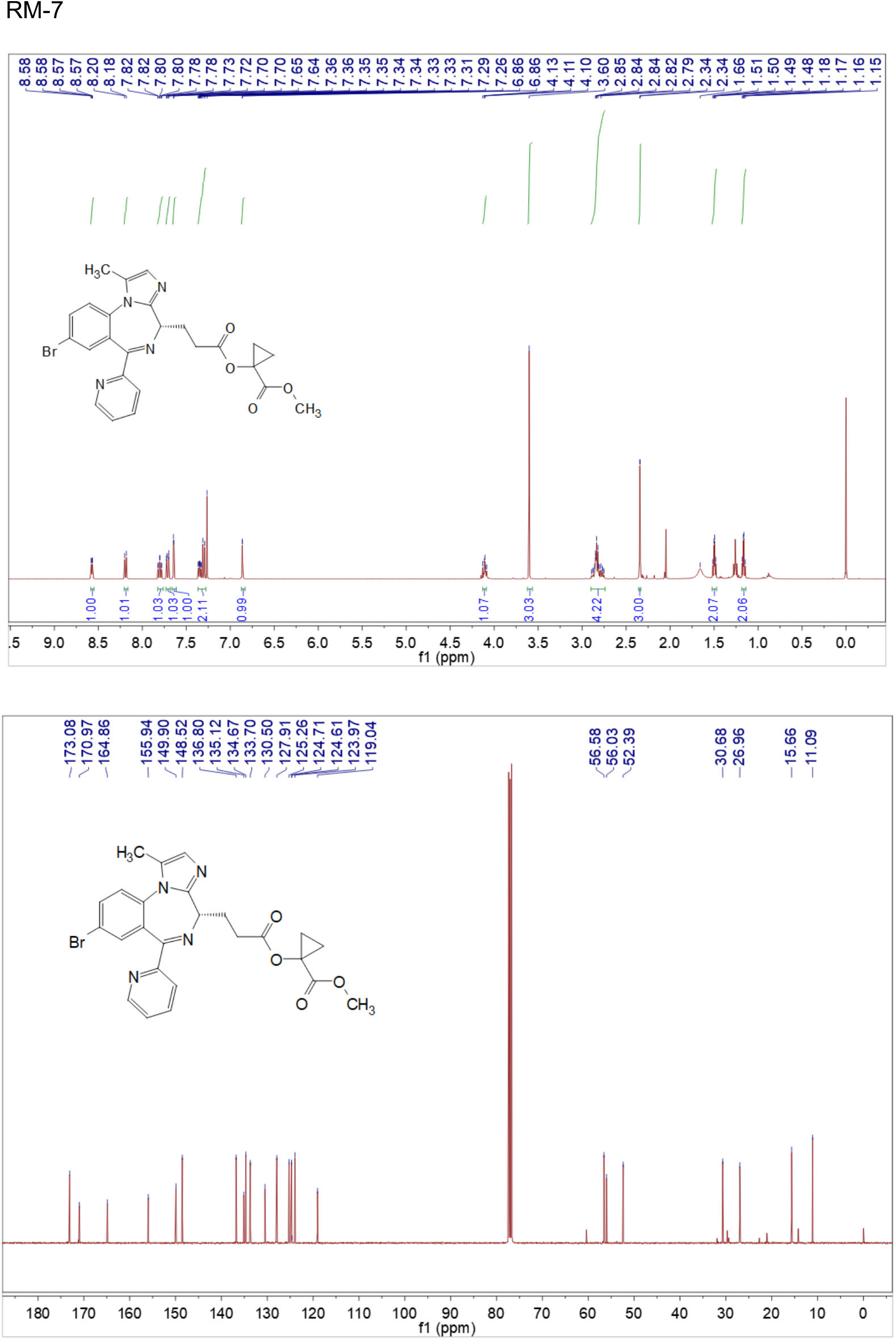

## Author Contributions

Z.X., W.A., Z.M., J.S., Z.W., W.L., Y.S., J.L., X.L., and B.K. designed research; Z.X., W.A. and Z.M. performed research; Z.X. contributed new reagents/analytic tools; Z.X. and W.A. analyzed data; and Z.X., W.A., X.L., and B.K. wrote the paper.

## Competing Interest Statement

The authors declare no competing interest.

## Notes

### Competing Interest Statement

The authors have declared no competing interest.

